# Non-viral vasculogenic reprogramming restores cognition and mitigates pathology in Alzheimer’s disease

**DOI:** 10.64898/2026.01.22.700900

**Authors:** Diego Alzate-Correa, Jonathan Stranan, Maria A. Rincon-Benavides, Natalia Areiza-Mazo, Luis F. Narvaez-Perez, Shara Jaramillo Garrido, Tracy Nguyen, Roma Patel, William R. Lawrence, Ludmila Diaz-Starokozheva, Ana I. Salazar-Puerta, Andrea Tran, Alex Valentine, Natalia C. Mendonca, Olivia Seline, Junyan Yu, Sydney Ripsky, McKayla Hagan, Stephen Piatkowski, Julie Fitzgerald, Fangli Zhao, Tatiana Z. Cuellar-Gaviria, Russell Lonser, Candice Askwith, Natalia Higuita-Castro, Olga Kokiko-Cochran, Daniel Gallego-Perez

## Abstract

Alzheimer’s Disease (AD) is characterized by progressive cognitive decline associated with amyloid-beta (Aβ) plaques, neurofibrillaiy tangles, inflammation, synaptic loss, and profuse neuronal death. Accumulating evidence demonstrates that cerebrovascular impairment precedes the emergence of neuropathological hallmarks, implicating vascular dysfunction as an early contributor to AD onset and progression. We investigated a non-viral strategy to generate pro-vasculogenic fibroblasts by transiently overexpressing *Etυ2, Foxc2,* and *Fli1 (EFF)* as a potential cell-based therapy for neurovascular deficits in AD. To assess therapeutic potential, ‘FFF-primcď fibroblasts were injected into a mouse model of AD (3xTg-AD) and wild-type controls via the intracerebroventricular (ICV) route, followed by cognitive assessments and subsequent brain tissue analyses. Our findings demonstrate that FFF-primed fibroblasts acquire vasculogenic properties, enhance cerebral blood flow (CBF), and alleviate spatial memory deficits in 3xTg-AD mice. Moreover, transplanted FFF-primed fibroblasts exhibited long-term survival, integrated into the brain vasculature, and promoted cortical vascular remodeling in the AD brain. Notably, ICV deployment of these cells is also correlated with reduced cortical amyloid-beta load, suggesting potential therapeutic benefits in reducing AD pathology. Transcriptomic analysis identified the activation of genes involved in fatty acid oxidation, such as Pparα, known for its anti-amyloidogenic and anti-inflammatory effects. Collectively, these findings highlight non-viral, reprogramming-based vasculogenic cell therapy as a promising strategy for Alzheimer’s disease, capable of alleviating cognitive decline and addressing AD pathology across cellular and tissue scales.

## Main

Alzheimer’s Disease (AD) is a form of dementia characterized by an accelerated decline in cognitive and executive capacities correlating with a substantial loss of synaptic connections and widespread neuronal death. Examination of AD affected brains also reveal two main neuropathological lesions, Aβ plaques and neurofibrillaiy tangles associated with chronic inflammatory process^1^. In addition, numerous studies indicate that impaired cerebrovascular function, including reduced Cerebral Blood Flow (CBF) and Blood-Brain Barrier (BBB) dysfunction, precedes the neuropathological changes, implicating these impairments with the onset and development of AD-type dementia^2–8^. Neuroimaging studies indicate that CBF deficits in AD patients occur well before the deposition of Aβ plaques^9^, and before brain atrophy or dementia can be detected^10^,^11^. Abnormalities in transport mechanisms and BBB breakdown seen in postmortem tissue^12^ further support a link between cerebrovascular dysfunction and AD and other dementias. Longitudinal studies in aging populations indicate that cerebral infarctions are a significant risk factor for dementia^13^,^14^. In addition, conditions that drive vascular pathologies, including hypertension, atherosclerosis, obesity, and diabetes, are significant risk factors for the development of AD^15–17^. Studies in animal models also reinforced this link. A recent study using a transgenic mouse model of Aβ plaque deposition showed that induction of cerebrovascular lesions by middle cerebral artery occlusion (MCAO) leads to Aβ accumulation around the infarct zone as a consequence of impaired Aβ clearance^18^. Another study showed that experimental induction of transient cerebral hypoperfusion in a mouse model of AD causes an exacerbation of the neuropathological lesions related to AD^19^. Additionally, different animal models carrying AD-related mutations showed a propensity to diminished CBF and BBB dysfunction^3^ ^4^, including mice with Amyloid Precursor Protein (APP) mutations *(e.g.,* Swedish, Florida, London), and mice with mutations in Presenilin-1, Tau, or Apoe4^20–31^. Importantly, recent studies show that increasing CBF by reducing capillary stalling in murine models of AD can lead to reduced disease burden^4–27^. Altogether, these studies suggest a strong link between cerebrovascular dysfunction and the development of AD-type dementias, and that therapeutic approaches aimed at correcting cerebrovascular deficits may delay and/or counteract neurodegeneration and functional decline.

Current therapies for AD are designed to target AD symptoms by enhancing glutamatergic and cholinergic neurotransmission in the remaining synapses^32^. Recently, Aβ-targeting immunotherapies have received regulatory approval and are being administered to select populations of AD patients^33^. Unfortunately, these treatments only manage to temporarily slow down the cognitive decline^34^, and despite extensive research and development, most of these agents have failed to show significant improvements in later stages of the disease. Consequently, there is an urgent need for new disease-modifying therapies to prevent, delay or reverse the symptoms of AD.

Stem cell-based therapies have emerged as a promising alternative strategy for the repair or replacement of deteriorating neurons and neuronal networks in AD^33–36^. Current strategies rely on the generation of new neurons from different sources including embryonic stem cells, mesenchymal stem cells, neural stem cells and induced pluripotent stem cells (iPSC). However, these stem cell-based therapies face numerous limitations. In many cases, progenitor or stem cell sources are insufficient, are difficult to isolate or pose major risks in terms of uncontrolled differentiation, tumorigenesis, ectopic integration into neural circuits, genetic abnormalities, and immunogenicity^37^. As an alternative to neuronal replacement, vasculogenic cell therapy offers a potential strategy to address cerebrovascular deficits in AD, leveraging the strong synergy between angiogenesis and neurogenesis during neural repair and development, wherein newly formed vasculature serves as a neurotrophic scaffold supporting neural regeneration.^38–40^. Pro-vasculogenic cell therapies are based on the generation and use of “exogenous” endothelial cells, or the stimulation of local/tissue-resident endothelial progenitor cells for therapeutic purposes^41^. Nevertheless, in diseases states like AD, stem/progenitor cell pools are extremely limited or difficult to isolate^42^. In addition, stem cell functionality can be compromised by the progression of AD, with amyloidosis and chronic inflammation impairing stem cell proliferation and differentiation capabilities^43^. Finally, the use of viral vectors to induce pluripotency of somatic cells poses major risks in terms of tumorigenesis, genetic abnormalities, and immunogenicity^44–48^. To overcome the major limitations of traditional stem/progenitor cell therapies, recent advances in direct nuclear reprogramming have opened up the possibility for the development of vasculogenic cell-based therapies utilizing more readily-available cell sources (*e.g.,* fibroblasts), and bypassing the need for induced pluripotency^49^. Here we used non-viral delivery of transcription factor genes *Etv2, Foxc2,* and *Fli1 (EFF)* into fibroblasts, to induce pro-vasculogenic/endothelial cell conversions (iECs)^50,51^ and study their therapeutic effect in a mouse model of AD. Our results show that intracerebroventricular deployment of EFF-primed fibroblasts leads to increased cerebral perfusion and vascularity, reduction of cognitive deficits, and reduction in Aβ accumulation.

## Results

### EFF electrotransfection rapidly initiates vasculogenic reprogramming in fibroblasts

Previously, we demonstrated that *EFF* delivery by electrotransfection led to vasculogenic reprogramming of fibroblasts into iECs within 1 weeks^52-55^ **(Fig. 1A).** To evaluate early transcriptome changes induced by the *EFF* cocktail, mouse fibroblasts were electrotransfected with expression plasmids encoding for the transcription factors *Etv2, Foxc2,* and *Fli1* at a 1:1:1 mass ratio. Sham/empty plasmids with the same backbone were used as controls. Successful transfection was confirmed by quantitative reverse transcription polymerase chain reaction (qRT-PCR) 24 hours post-deliveiy, showing robust overexpression of *EFF* compared to the sham/control group **(Supp. Fig. 1A).** To profile transcriptional changes, we conducted RNA sequencing (RNA-seq) on mRNA from naïve fibroblasts *(i.e.,* no electrotransfection) and from fibroblasts 24 hours after electrotransfection with either *EFF* or sham plasmids. Principal Component Analysis (PCA) showed strong intra-group clustering and distinct inter-group segregation, indicating robust transcriptional differences between conditions **(Supp. Fig. 1B).** Subsequent pairwise comparison between the three experimental groups using thresholds of Log2FC> 111 and false discovery rate (FDR)-adjusted p-value (q-val) <0.05 also illustrate marked differences in gene expression caused by the *EFF* cocktail. Comparison between sham versus naïve cells identified 1687 significantly differentially expressed genes (DEGs), with 956 genes downregulated and 731 genes upregulated in the sham group relative to the naïve group **(Fig. 1B, Supp. Fig 1C and Supp. Table 1).** Further analysis of the DEGs list by Gene ontology (GO) uncovered an increased representation of genes associated with the regulation of immune responses, represented in GO categories that include *leukocyte migration* (94 DEGs), *positive regulation of cytokine production* (103 DEGs), *regulation of leukocyte activation* (104 DEGs), *adaptive immune response* (99 DEGs), *response to molecule of bacterial origin* (90 DEGs) and *T-cell activation* (96DEGs) among other similar categories. In addition, there was also a significant representation of DEGs involved in *positive regulation of cell migration* (90 DEGs), and DEGs related to *angiogenesis* (89 DEGs) **(Fig. 1C).** However, from these 89 DEGs associated with angiogenesis, 58 were also present in GO categories related with immune responses or cell migration such as *Syk, Ccr2, Vegfc , Itgb2, Ackr3, Ccr5, Ccl5, Cybb, Celli* and *Mmpg* **(Fig. 1D,E),** whereas 31 are exclusively related with angiogenesis including *Notch4, Angptl6, Vash1, Optc, Rspo3, Angptl4* and *Cysltr1* **(Fig. 1F).** Furthermore, analysis by Gene Set Enrichment Analysis (GSEA)^56–57^ operating with *Hallmark Gene Sets* (with FDR q-val <0.25) indicate that *Angiogenesis* have a normalized enrichment score (NES) of 1.58, a number below other hallmarks such as *Interferon gamma response, Apoptosis, and TNFa Signaling via NFKB,* with NES of 1.93, 1.91 and 1.81 respectively **(Fig. 1G and Supp. Table 2).** According to these results, there are substantial changes in gene expression in fibroblasts induced simply by the electrotransfection procedure.

**Figure 1.**
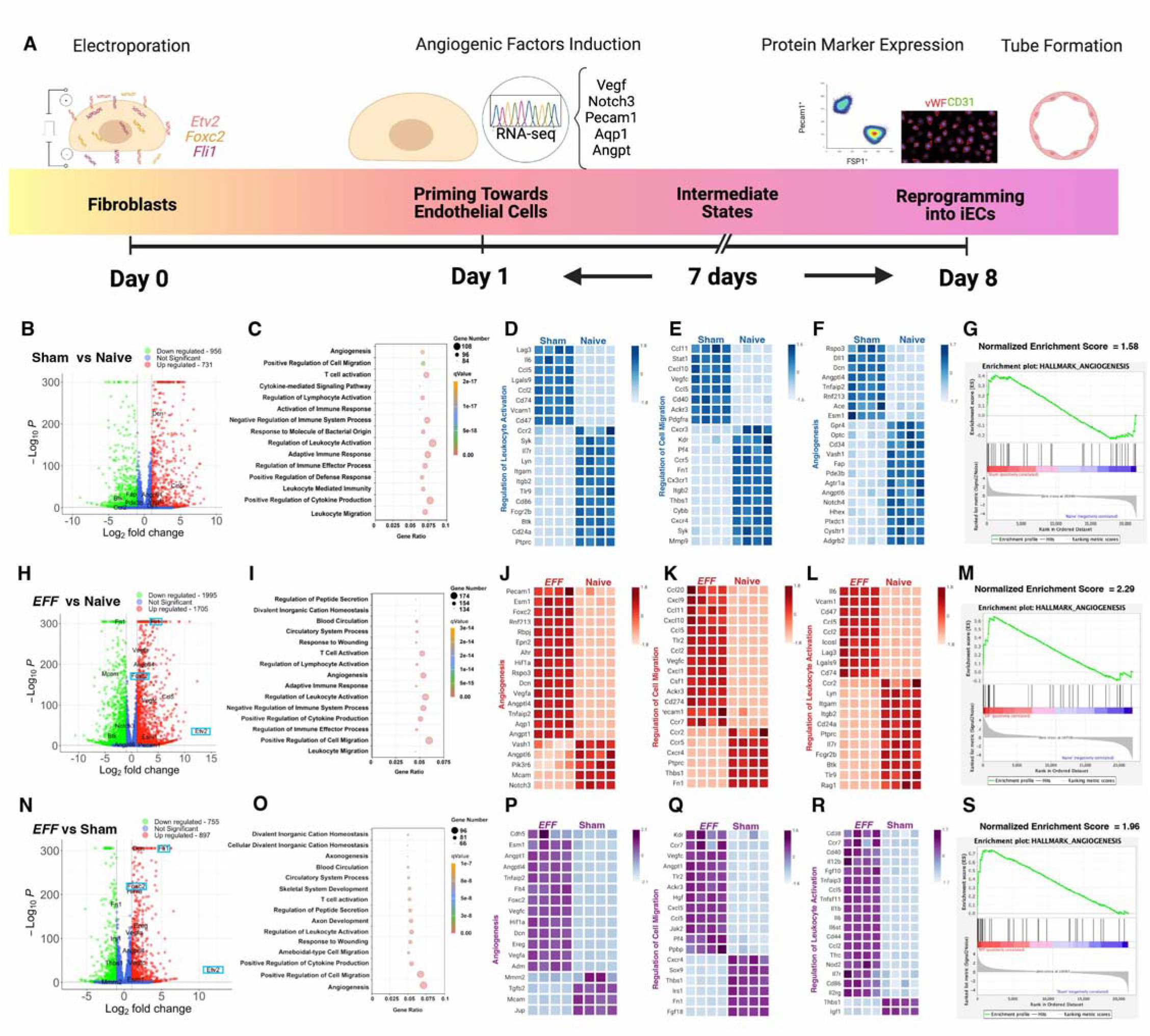
Electrotransfection of fibroblasts with the transcription factors *Etv2, Foxc2,* and *Flii (EFF)* induces an early vasculogenic priming. **(A)** Progression of induced cell differentiation after electrotransfection with the *EFF* factors showing early transcriptional changes, together with the expression of endothelial markers and endothelial functions by day 8. RNA-seq analysis of fibroblasts 24h after electrotransfection with a sham plasmid or the FFF-cocktail (n=4). As an additional control, non-electrotransfected fibroblasts were included (Naïve) (n=4). (B) Volcano plot illustrating DEGs between naive and fibroblasts electrotransfected with a sham plasmid (Log_2_FC > 1: adjusted p-value ≤ 0.05). (C) Gene Ontology (GO) analysis with the biological processes enriched in the lists of DEGs. Heatmaps representing the relative expression differences in key DEGs for the main GO biological process including **(D)** regulation of leukocyte activation, **(E)** regulation of cell migration and **(F)** angiogenesis. **(G)** GSEA analysis representing hallmark gene sets highlights angiogenesis as an enriched biological process in our transcriptome evaluation. Analysis under the same parameters were performed to compare Naive cells with fibroblasts electrotransfected with the *EFF* plasmid cocktail and represented with a **(H)** volcano plot, **(I)** GO enrichment, heatmaps for key DEGs on **(J)** angiogenesis, **(K)** regulation of cell migration and **(L)** regulation of leukocyte activation. **(M)** GSEA representation of angiogenesis hallmark. Finally, results of comparison between cells electrotransfected with a sham plasmid compared to the cells electrotransfected with the *EFF* plasmid cocktail are represented by a **(N)** volcano plot, **(O)** GO enrichment, heatmaps for key DEGs on **(P)** angiogenesis, **(Q)** regulation of cell migration and (R)regulation of leukocyte activation. **(S)** GSEA representation of angiogenesis hallmark.

Subsequent comparison between EFF-transfected cells against naïve cells identified larger changes in gene expression, with a total of 3700 DEGs comprising 1995 downregulated genes and 1705 upregulated genes in the *EFF* group when compared with the naïve group **(Fig. 1H, Supp. Fig. 1D and Supp. Table 3).** Of note, *Etv2, Foxc2* and *Fli1* were significantly upregulated in EFF-transfected cells validating our electrotransfection protocol **(Fig. 1H).** GO analysis in this case revealed significant representation of DEGs related to *angiogenesis* (157 DEGs), *circulatory systems process* (137 DEGs) and *blood circulation* (136 DEGs), together with *positive regulation of cell migration* (174 DEGs). In addition to these, GO revealed the regulation of immune process with similar GO categories to those identified when comparing sham versus naïve cells, including *leukocyte migration* (134 DEGs), *positive regulation of cytokine production* (148 DEGs), *regulation of leukocyte activation* (163 DEGs), *adaptive immune response* (135 DEGs), and *T-cell activation* (153 DEGs) **(Fig. 11).** In this case, from the 157 DEGs related with *angiogenesis,* 89 were also related with immune processes *(Ccl2, Ccl5, Vegfc, Ackr3, Pcami, Fni, Ccr2, Itgb2)* **(Fig. 1K, L),** and 68 are exclusively related with *angiogenesis (Notch3, Esmi, Rnf2i3, Rspo3, Vash1, Vash2, Angptl4, Angptl6)* **(Fig. 1J).** However, if we pool DEGs from similar GO categories like *angiogenesis, circulatory systems process,* and *blood circulation,* we found a total of 348 unique DEGs, out of which 153 are shared with immune GO categories, and 195 DEGs are exclusive to angiogenic-related GO categories. Additional analysis by GSEA also indicates a strong enrichment of *Angiogenesis* genes with a NES of 2.29, whereas *Interferon gamma response, Apoptosis* and *TNFa Signaling via NFKB* show lower NES of 1.89, 2.15 and 2.08 respectively **(Fig.1 M and Supp. Table 4).** These results show an early (24h post-electrotransfection) and robust induction of vasculogenic transcription program, initiated by the induced expression of the *EFF* reprogramming cocktail.

Finally, comparison between EFF-transfected and Sham-transfected cells identified 1652 DEGs, 755 downregulated, and 897 upregulated in the *EFF* groups with reference to the sham group, including again an upregulation of our transfected genes *Etv2, Foxc2 andFli1* **(Fig. 1N. Supp. Fig. 1E and Supp. Table 5).** In this case, GO analysis indicated again a strong enrichment of genes associated with *angiogenesis* (96 DEGs), *circulatory system process* (68 DEGs), *blood circulation* (67DEGs) and regulation of vasculature development (65 DEGs) **(Fig. 1O),** further corroborating our previous results, indicating a transcriptional shift from a fibroblast to an endothelial phenotype, and reinforcing our observation that vasculogenic reprogramming is initiated within the first few hours after *EFF* electrotransfection. Immune related GO categories represented in this comparison include *regulation of leukocyte activation* (74DEGs) and *positive regulation of cytokine production* (74 DEGs) **(Fig. 1O).** Surprisingly, our analysis also shows a representation of *axon development* (73DEGs) and *axonogenesis* (66 DEGs). Detailed analysis of these GO categories shows a total of 143 unique DEGs included in our angiogenic GO categories, from which 30 DEGs are shared with immune related GO categories *(Cd38, Jak2, Hgf Vegfc, Ccl5)* **(Fig. 1Q, R),** and 113 DEGs are exclusive to angiogenic categories *(Cdh5, Adm, Nos2, Jup, Foxc2, Angptl4)* **(Fig. 1P).** GSEA analysis with our DEGs indicates a NES of 1.96 for *Angiogenesis, Inflammatory response, Apoptosis, and TNFa Signaling via NFKB* have NES values of 1.93, 1.96 and 1.80 respectively **(Fig 1S and Supp. Table 6).** Taken together, our analysis of gene expression shows that 24 hours after electrotransfection there is an early and robust expression of our reprogramming factors *Etv2, Foxc2,* and *Fli1,* that is accompanied by transcriptional changes fitting an induced vasculogenic commitment of our cells. In parallel, we also observed a significant activation of immune-related processes in our electro transfected cells, as well as GO and GSEA categories suggestive of a synergy between angiogenesis and inflammation, including *positive regulation of cell migration, response to wounding, and* Hypoxia **(Fig. 1C, I, O).** Together, these findings indicate that EFF-electrotransfected fibroblasts mount a robust pro-angiogenic response, as early as 24 hours, as evidenced by the upregulation of key vasculogenic mediators such as *Vegfa* and *Vegfcm* **(Supp. Tables 3 and 5),** which can drive endothelial proliferation, migration, and tube formation. In addition, *EFF* reprogramming induced differential expression of genes involved in axon development, including several *Semaphorin* family members and *Reelin,* suggesting potential effects on neural plasticity and synaptic remodeling.

Beyond the early pro-vasculogenic transcriptional changes observed after *EFF* electrotransfection, we assessed whether EFF-transfected fibroblasts also release extracellular vesicles (EVs) cariying the *EFF* factors. Using a microplate insert system, and consistent with our previous findings^51–55^, we detected transfer and uptake of EFF-loaded EVs by neighboring fibroblasts **(Supp. Fig. 2),** indicating a paracrine mechanism capable of propagating *EFF* delivery and potentially amplifying vasculogenic reprogramming in adjacent cells. Such EV-mediated signaling suggests that, upon transplantation, therapeutic effects may arise not only from the injected cells themselves but also from secondary reprogramming induced in surrounding host cells. The release of EVs carrying *EFF* factors provides a complementary paracrine mechanism capable of transferring vasculogenic cues to neighboring cells, thereby amplifying the overall angiogenic and vasculogenic impact beyond the directly transfected population.

**Figure 2.**
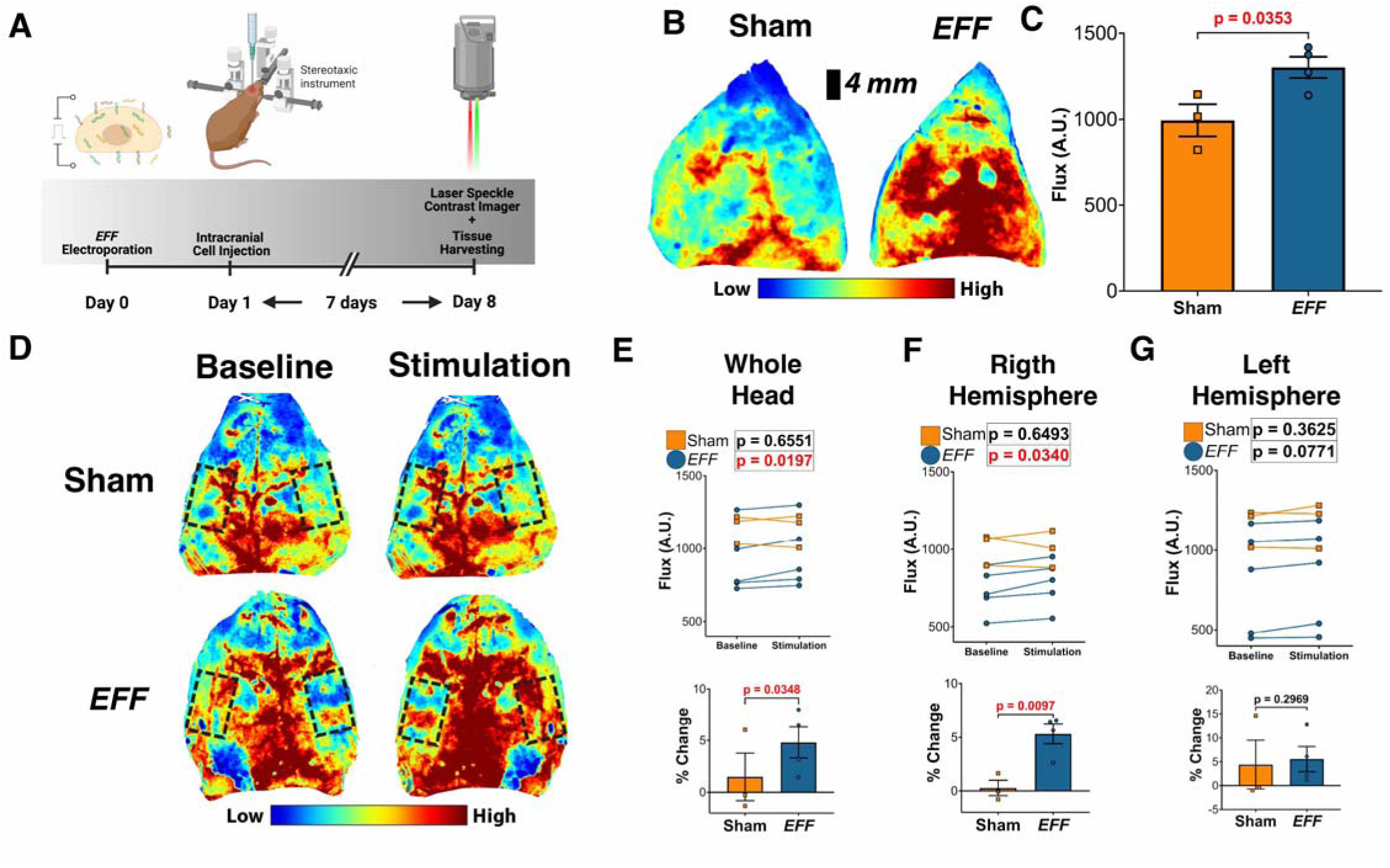
Intracranial injection of EFF-electrotransfected fibroblasts induces an increase in cerebral blood flow and increases neurovascular coupling. **(A)** Illustration of the experimental procedure followed to test changes in CBF after intracranial injection of cells. Fibroblasts were pre-labeled with BrdU for 48h followed by electroporation with sham or *EFF* plasmids. 24h after electrotransfection, cells were intracranially injected in 6-month-old 3xTg-AD mice. Seven days after injection, global CBF was measured using laser speckle imaging (LSI). **(B)** Representative images of relative perfusion obtained by LSI on the skull of 3xTg-AD mice injected with Sham- or EFF-electrotransfected cells. **(C)** Quantification of perfusion by LSI indicates an increase in global CBF after injection of FFF-transfected cells relative to animals injected with sham plasmid electroporated cells (n=3-4. Student t-test). (D) For neurovascular coupling analysis, 7 days after intracranial injection of electrotransfected fibroblasts mice were subjected to LSI recordings while mechanically stimulating their whiskers. Representative images of 3xTg-AD mice injected with sham cells (top) or FFF-primed cells (bottom) during the pre-stimulation baseline recordings (left) and during the stimulation period (right). **(E)** LSI recordings of the entire head during baseline and stimulation for each mouse (top) and quantification of the percentage change for sham andEFF(n=3-5) experimental groups (bottom). **(F)** Similar quantification was performed in the right hemisphere while stimulating the left whiskers and (G) in the left hemisphere while stimulating the right whiskers (n=3-5. Student t-test)

### EFF-primedfibroblast transplantation enhances cortical cerebral bloodfloiv in AD mice

Previous studies reported that intracranial transplantation of EFF-primed fibroblasts enhances cerebral blood flow (CBF) under both physiological and ischemic conditions^51^. To understand the effect of EFF-primed fibroblasts on CBF in the context of AD, we stereotaxically injected sham- or EFF-primed fibroblast in the lateral ventricles (LVs) of 6-month-old 3xTg-AD mice and evaluated cortical CBF by laser speckle imaging (LSI) **(Fig. 2A).** We delivered the cells into the LVs to promote widespread distribution across the brain, as this route provides direct access to the cerebrospinal fluid (CSF) and brain cisterns^58^. Once the CSF reaches the fourth ventricle, it circulates through the subarachnoid space surrounding the brain and interfaces directly with the cerebral vasculature via the Virchow-Robin and perivascular spaces^59^. This deliveiy route leverages the CSF for brain-wide cell distribution, resembling the intrathecal administration of cells used in clinical studies ^60–63^. Analysis of cortical CBF by LSI indicates that EFF-primed fibroblasts induced an increase in basal CBF when compared with animals injected with sham-transfected cells **(Fig. 2B, C).**

To further assess the effects of EFF-primed cells on CBF, we employed a similar injection schedule and quantified CBF at rest and during bilateral whisker stimulation. This paradigm elicits a sensoĩy-evoked CBF response in the barrel cortex, which reflects functional neurovascular coupling^64–65^, a mechanism previously reported to be impaired in the 3xTg-AD model^66^. In this case, 7 days after stereotaxic injection of the cells in the LVs, LSI recording were taken for one minute under baseline conditions followed by 30s stimulation of the right-side whiskers, one minute of basal recordings and 30s stimulation of the left side whiskers. This entire protocol was applied twice. Whole-brain CBF analysis revealed that injection of EFF-primed cells enhanced the perfusion response to whisker stimulation compared to baseline **(Fig. 2D).** This was evident from the increased difference in perfusion between stimulation and baseline periods **(Fig. 2E top),** as well as from a greater percentage change relative to sham-transfected controls **(Fig. 2E bottom).** Further analysis of right-hemisphere responses induced by left whisker stimulation revealed significant differences in perfusion levels in mice injected with EFF-transfected cells when comparing baseline and stimulation conditions **(Fig. 2D, F).** Surprisingly, stimulation of the right whiskers did not elicit significant differences in perfusion on the left hemisphere **(Fig. 2D, G),** which may reflect hemispheric variability in cerebrovascular reactivity and/or treatment distribution. Together, these results validate our prior observations that EFF-primed fibroblasts enhance cortical perfusion under both physiological and ischemic conditions and further suggest a beneficial effect on neurovascular coupling.

### High-dose EFF-primed fibroblast delivery improves cognition in AD mice

Having established the ability of EFF-primed fibroblasts to enhance cerebral perfusion and support neurovascular coupling, we next examined whether they can modulate cognitive performance in 3xTg-AD mice **(Fig. 3).** Wild-type (WT) mice receiving the same treatment served as controls. To overcome the cell number limitations inherent to intracranial delivery, we employed a high-dose, three-injection regimen in which fibroblasts were electro transfected with *EFF* or sham plasmids and injected 24 hours later into the LVs of 3xTg-AD and WT mice. Injections were spaced one month apart (at 16, 20, and 24 weeks of age) to permit recovery and maximize cumulative cell delivery **(Fig. 3A).** This dosing strategy was motivated by prior results showing that a single EFF-fibroblast injection improved basal and sensory-evoked CBF, while the absence of left-hemisphere responsiveness indicated that higher cell doses might be necessary for complete neurovascular restoration. Furthermore, this three-dose regimen parallels clinical ICV cell delivery protocols for AD, where repeated monthly administrations were shown to be safe and feasible^67^. Thus, our repeated delivery strategy maintains translational relevance by mirroring clinically tested dosing intervals and cumulative exposure. Of note, for the final injection fibroblasts were pre-labeled with 5-Bromo-2’-Deoxyrrridine (BrdU) 48h before electrotransfection to facilitate long-term tracking of the injected cells. Two weeks after the last injection, we assessed spatial learning and memory using the Barnes Maze, and evaluated recognition memory with the Novel Object Location (NOL) and Novel Object Recognition (NOR) tasks. All these are hippocampus-dependent tasks and have been shown to be impaired in 3xTg-AD mice^68–69^

**Figure 3.**
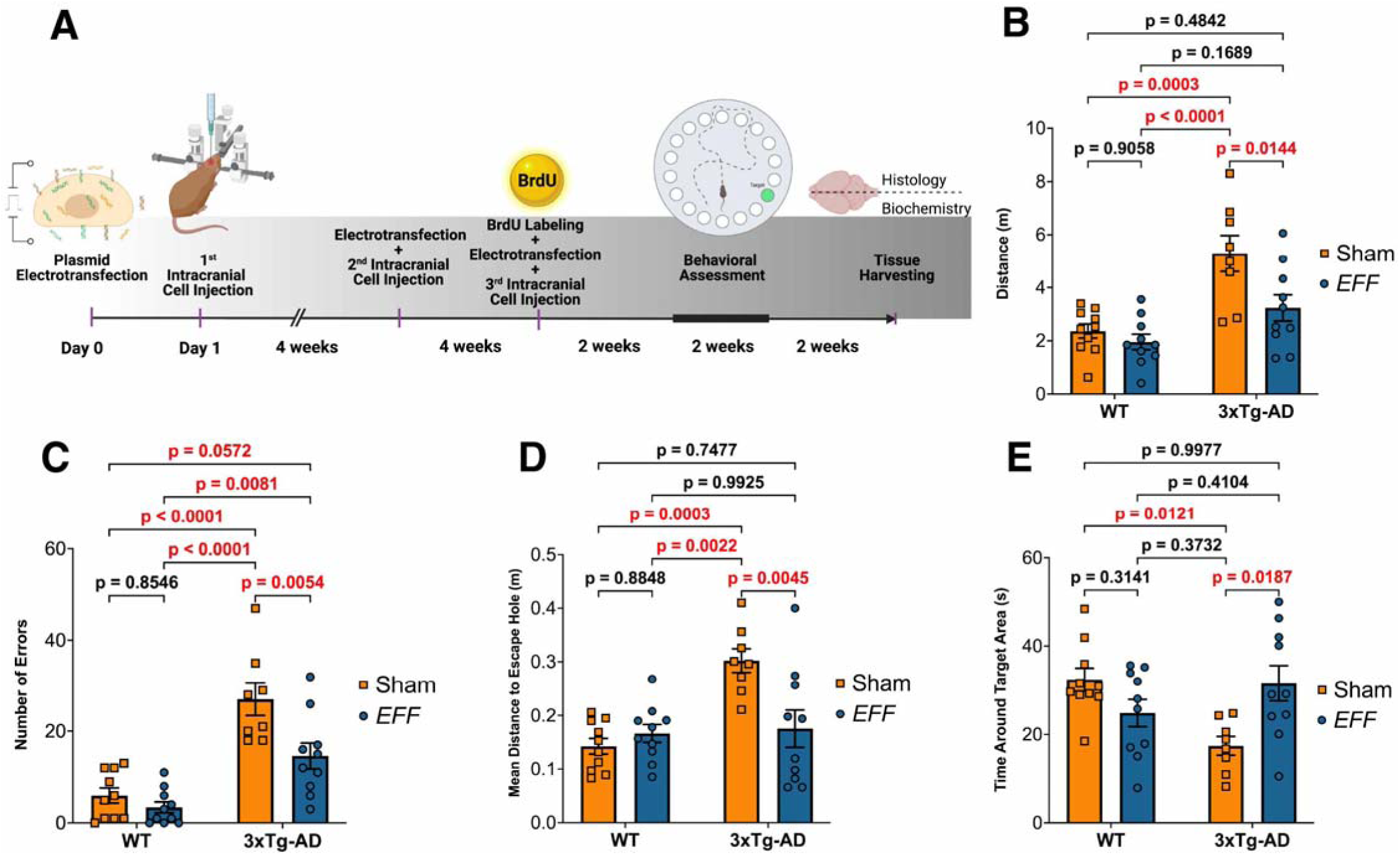
Serial injections of fibroblasts electrotransfected with *EFF* reduce cognitive capacity impairments in 3xTg-AD mice. **(A).** Illustration of the experimental timeline describing the monthly electrotransfection and intracranial injections of fibroblasts in WT and 3xTg-AD mice. All the animals receive 3 bilateral injections of 4x104 cells. Two weeks after the last injection, recognition and spatial memory was evaluated. Two weeks later brain tissue was harvested. Evaluation of spatial memory with the Barnes maze. Twenty-four hours after the end of the acquisition phase, retrieval of the task is evaluated by removing the escape box and quantifying the retention of the escape box location. Parameters evaluated include **(B)** distance traveled, **(C)** number of errors, **(D)** mean distance from the target hole and (E) time spent in proximity of the target hole (n= 8-10, Two-way ANOVA).

During the Barnes Maze acquisition phase, animals performed three 2-minrrte trials per day for five days, using distal spatial cues to locate a concealed escape box. Multiple parameters were analyzed to assess learning **(Supp. Fig. 3A-F).** As expected and consistent with prior reports^70^, genotype differences dominated acquisition performance, with 3xTg-AD mice showing poorer learning than WT controls. However, no significant differences were observed between EFF-primed and sham cell-treated groups within each genotype. Learning progression metrics, including the acquisition index and savings index, were comparable across all treatment conditions **(Supp. Fig. 3G, H).** In contrast, group differences emerged during the retrieval probe conducted 24 hours after the final acquisition trial, when the escape box was removed to assess memory for its previous location **(Fig. 3B-E).** Here, 3xTg-AD mice injected with sham cells exhibited clear impairments, traveling longer distances, committing more errors, and maintaining a greater mean distance from the target hole compared with WT animals. Notably, 3xTg-AD mice treated with EFF-primed fibroblasts showed marked improvements across these measures, traveling less distance **(Fig. 3B),** making fewer errors **(Fig. 3C),** and displaying reduced mean distance to the target location **(Fig. 3D),** reaching values comparable to WT controls. Time spent in the target hole area showed a similar rescue, with the 3xTg-AD-EFF group exhibiting increased target engagement relative to 3xTg-AD-Sham mice, again normalizing to WT levels **(Fig. 3E).** Additional retrieval metrics, including time and path in the target quadrant, as well as latency to reach the target area, did not differ significantly across groups **(Supp. Fig. 3I-K).**

To complement the Barnes Maze results, we evaluated recognition memory using the NOL and NOR tasks. In both assays, recognition indices were similar between 3xTg-AD and WT mice, and EFF-primed cells did not alter performance in either genotype **(Supp. Fig. 3L,M).** Total exploration times were comparable across groups **(Supp. Fig. 3N,P).** For the NOL task, only 3xTg-AD mice receiving sham cells showed a shift toward exploring the non-relocated object **(Supp. Fig. 3O),** suggesting a mild deficit relative to the other groups. In the NOR task, WT mice, but not 3xTg-AD mice, demonstrated the expected preference for the novel object^71^, with no effect of *EFF* treatment **(Supp. Fig. 3Q).** Overall, recognition memory measures did not show treatment-related improvements, underscoring that the most robust cognitive benefits of EFF-primed cells emerged in the Barnes Maze spatial memory test. To evaluate whether the EFF-associated spatial memory benefits reflected changes in synaptic plasticity in 3xTg-AD mice, we conducted electrophysiological analyses; however, no differences were detected between sham and *EFF* groups **(Supp. Fig. 4).**

**Figure 4.**
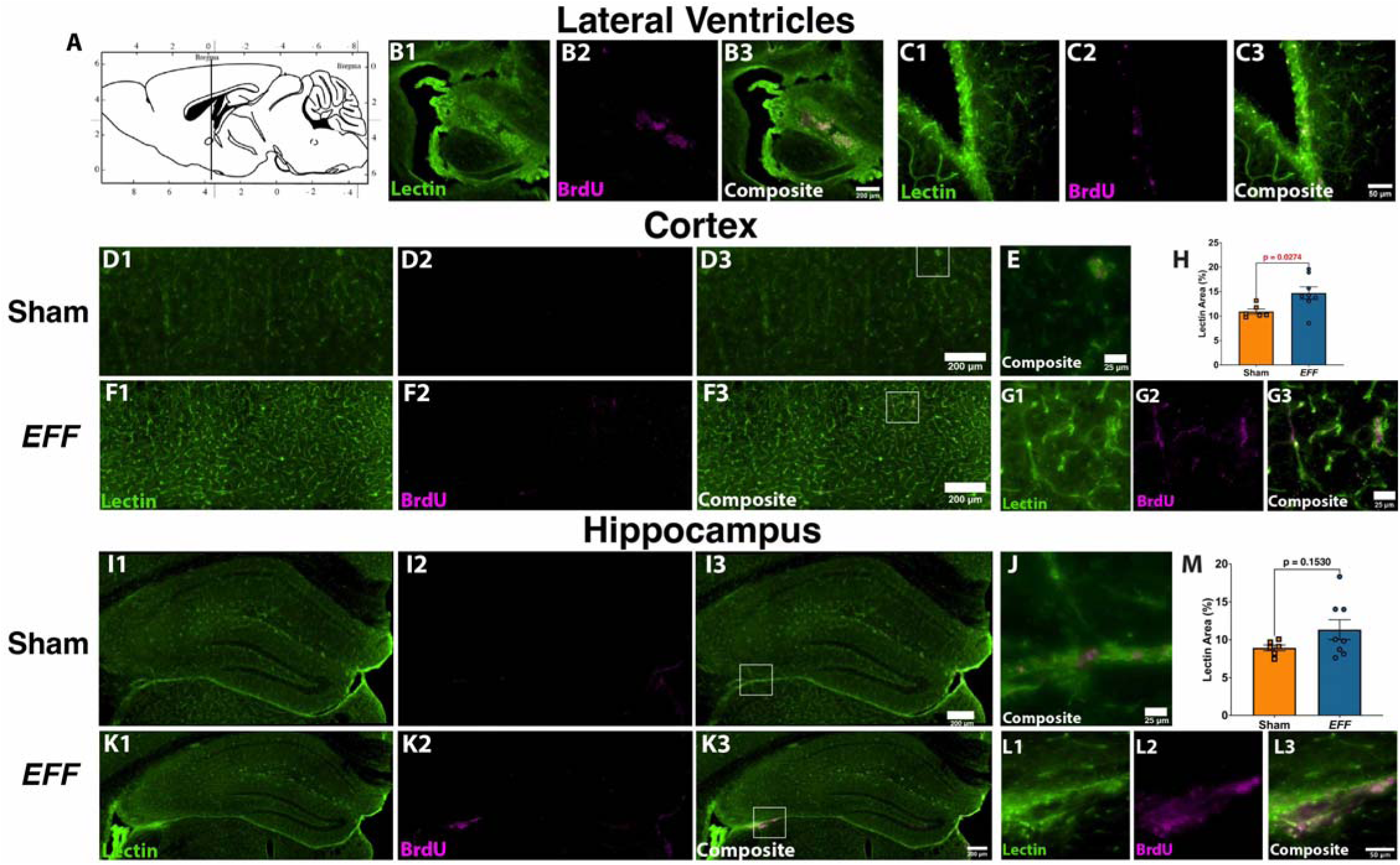
Intracranially injected EFF-primed cells can migrate beyond the injection site, survive long-term and increase vascularity in 3xTg-AD mice. **(A)** Sagittal illustration from the mouse brain atlas depicting the injection target site in the LVs. Immunofluorescence microphotographs of staining against BrdU (magenta) and blood vessels using lectin (green) in brain sections from mice 6 weeks after the last injection of EFF-primed cells. Fibroblasts were pre-labeled with BrdU, allowing specific tracking of the cells after injection and their position relative to blood vessels. **(B1-3)** Microphotograph of the LV showing the generation of a cell scar in the injection site of a 3xTg-AD mice injected with EFF-primed fibroblasts. **(C1-3)** Microphotograph of the LVs ependymal layer also showing the presence of BrdU positive cells. **(D1-3)** Low magnification photograph of the somatosensory cortex of 3xTg-AD mice injected with sham-electrotransfected cells. A higher magnification of the area depicted in **D3** is shown in **E. (F1-3)** Low magnification photograph of the somatosensory cortex of a 3xTg-AD mouse injected with EFF-primed cells. A higher magnification of the area depicted in **F3** showing BrdU positive cells in close contact with blood vessels is showed in **G1-3. (H)** Quantification of somatosensory cortex lectin shows an increase in the vascular area in the 3xTg-AD mice treated with EFF-transfected cells (n=6-8. Student t-test). **(I-J)** Similar observations were made in low and high magnification images of the hippocampus of 3xTg-AD mice injected with sham-electrotransfected cells, and **(K-L)** 3xTg-AD mice injected with EFF-primed cells. (M) Quantification of lectin-positive area in the hippocampus shows no differences between groups (n=6-8. Student t-test).

Altogether, our results indicate that EFF-primed fibroblasts exert a positive effect on the cognitive performance of 3xTg-AD mice, with the most robust improvements observed in the Barnes Maze spatial memory test. In contrast, recognition memory tasks (NOL and NOR) did not reveal treatment-related benefits, and in some cases showed minimal to no genotype differences, suggesting that at this age 3xTg-AD mice may exhibit milder deficits that are less amenable to detection and/or rescue by these assays. These findings underscore that the cognitive enhancement produced by EFF-primed cells is most evident in domains with clearer phenotype expression, supporting the therapeutic potential of EFF-primed fibroblasts to improve spatial learning and memory in AD.

### EFF-primed fibroblast therapy improves cortical vascularization in AD mice

Two weeks after behavioral evaluation, brains from all the experimental groups were harvested and cut along the midline to separate both hemispheres. One hemisphere was further dissected to isolate the hippocampus, striatum, cerebellum and cortical tissue for biochemical analysis. The other hemisphere was preserved for immunohistochemical (IHC) analysis by fixation and cŗyopreservation. Tissue reserved for IHC was cŗyo- sectioned to obtain serial free-floating sections. To analyze the vascular area and track the biodistribution of our injected cells, we performed a double fluorescence staining using *Lycopersicon Esculentum* Lectin (LEL) fused to the fluorophore DyLight 488 to detect brain blood vessels, together with indirect immunofluorescence staining targeting BrdU with secondary antibodies conjugated to the fluorophore Alexa 647. Initial examination of brain sections encompassing the LVs from 3xTg-AD mice injected with EFF-primed fibroblasts revealed the formation of highly vascularized tissue surrounding BrdU^+^ signal, as well as BrdU labeling within the ependymal layer of the ventricular zone **(Fig. 4A-C),** suggesting successful targeting and engraftment within the LVs. Similarly, in 3xTg-AD mice injected with either sham- or EFF-primed fibroblasts, we detected injected cells within cortical regions well beyond the injection site, including the somatosensory cortex. In these areas, BrdU^+^ cells were frequently observed in close proximity to lectin^+^ vasculature, suggesting an association between the transplanted cells and blood vessels **(Fig 4D-G).** We also detected BrdU^+^ cells within the hippocampal region **(Fig. 4I-L).** Some of these cells were located in areas of the hippocampus adjacent to the brain cisterns **(Fig. 4J, L),** as well as within the hippocampal parenchyma. Quantification of cortical vascular area revealed that 3xTg-AD mice injected with EFF-primed fibroblasts exhibited a significant increase in lectin^+^ area compared with those injected with sham-transfected fibroblasts **(Fig. 4H).**

In contrast, no significant differences in vascular area were observed in the hippocampus **(Fig. 4M).** Parallel analyses in WT mice also revealed BrdU^+^ cells in the somatosensory cortex **(Supp. Fig. 5A, B)** and hippocampus **(Supp. Fig. 5D, E).** However, no differences in lectin^+^ area were detected in these regions across the sham and *EFF* groups **(Supp. Fig. 5C, F).**

**Figure 5.**
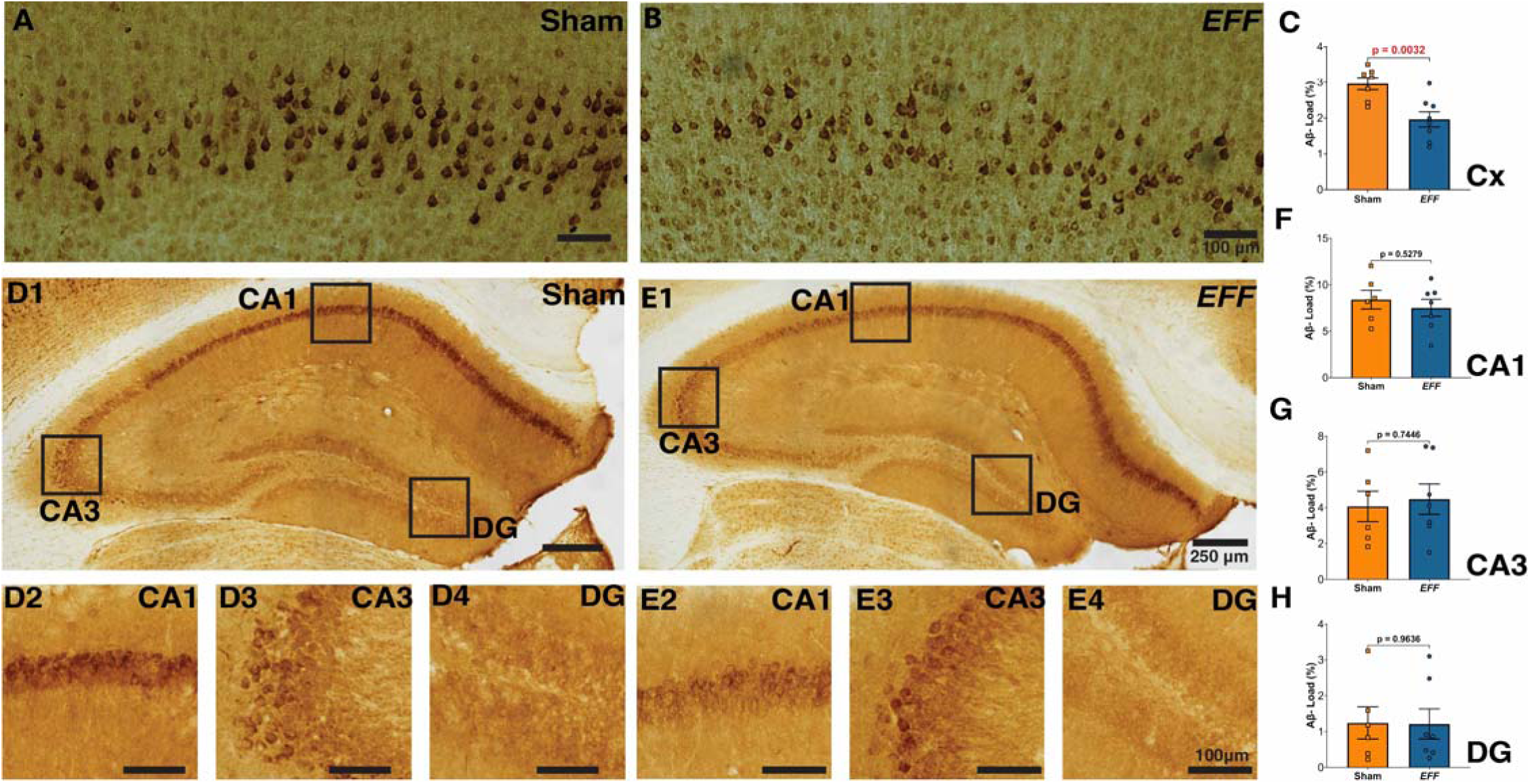
Injections of fibroblast electrotransfected with the *EFF* factors reduce intracellular accumulation of amyloid beta in 3xTg-AD mice. Immunohistochemical staining against Aβ in the somatosensory cortex of 3xTg-AD mice injected with sham- **(A)** or EFF-primed cells **(B). (C)** Quantification of Aβ in the cortex indicated a reduction in the load in mice injected with EFF-transfected cells. Amyloid beta immunohistochemistry in the hippocampus of 3xTg-AD mice injected with sham- **(D)** or EFF-primed cells **(E).** Inserts show higher magnification images of the principal cell layers in the CAi, CA3 and DG subregions. Quantification of Aβ load in the hippocampus subregions CAi **(F),** CA3 **(G)** and DG **(H)** show no differences in Aβ levels (n= 8-10. Student’s t-test).

Together, these analyses demonstrate that EFF-primed fibroblasts survive in the brain parenchyma for at least six weeks following injection and can migrate to regions distal to the delivery site. Importantly, these cells localize near endogenous blood vessels and elicit a measurable increase in vascular area within the somatosensory cortex of 3xTg-AD mice, underscoring a sustained pro-vasculogenic effect *in vivo*.

### EFF-primed fibroblast therapy reduces amyloid-β load in AD mice

To assess amyloid-β (Aβ) levels in 3xTg-AD mice, we performed IHC using a biotin-conjugated mouse monoclonal antibody against the Aβ peptide, followed by avidin-peroxidase detection with 3,3’- diaminobenzidine (DAB). Aβ immunoreactivity was detected in both the cortex and hippocampus. In the somatosensory cortex, Aβ labeling displayed the expected distribution across layers IV and V^70^ **(Fig. 5A,B).** Quantification revealed a significant reduction in cortical Aβ load in mice treated with EFF-primed fibroblasts compared with sham-treated controls **(Fig. 5C),** consistent with the increased vascular area observed in this region. In contrast, no differences in Aβ levels were detected in the hippocampus **(Fig. 5D-H),** indicating a region-specific effect of the *EFF* treatment. Additional histological staining for the assessment of neuroinflammation revealed no detectable changes associated with *EFF* treatment **(Supp. Figs. 6).**

**Figure 6.**
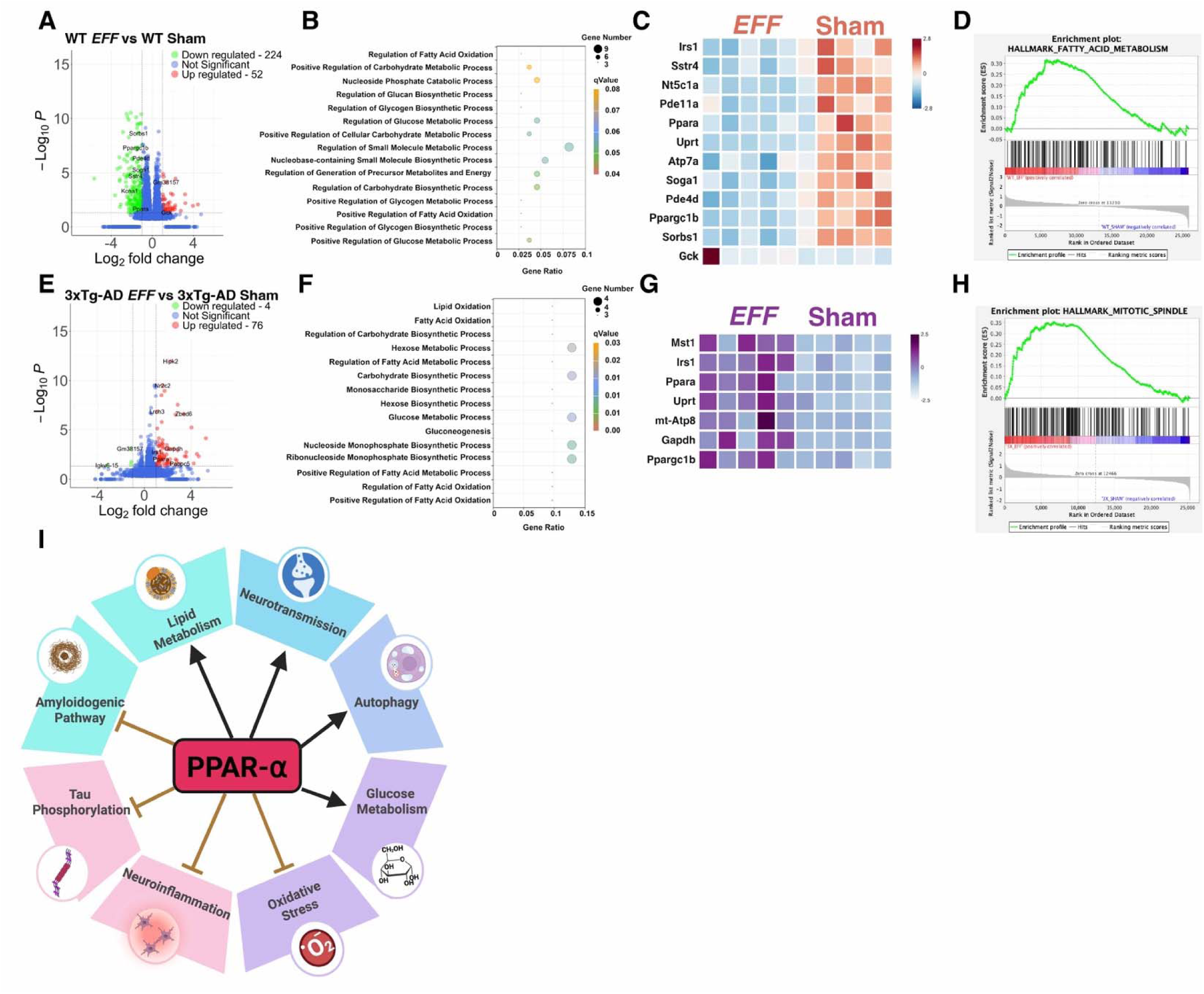
Transcriptome changes in the somatosensory cortex of WT and 3xTg-AD mice after injections of fibroblast electrotransfected with the *EFF* factors. RNA-seq analysis of somatosensory cortex tissue obtained 6 weeks after the last injection of fibroblasts electrotransfected with sham plasmid or the *EFF* factors in WT and 3xTg-AD mice (n=5). (A) Volcano plot illustrating DEGs in WT mice injected with EFF-primed fibroblasts contrasted with WT mice injected with sham electrotransfected cells. **(B)** GO analysis results indicating the main biological processes enriched in the DEGs list. **(C)** Heatmap illustrating the relative expression of 12 key genes related to the main GO categories. **(D)** GSEA analysis representing hallmark gene sets highlight fatty acid metabolism as the main enriched biological process. Results from analysis of 3xTg-AD mice injected with EFF-primed fibroblasts contrasted with 3xTg-AD mice injected with sham electrotransfected fibroblasts are represented by **(E)** a volcano plot, **(F)** GO of biological processes, and **(G)** heatmap clustering depicting the expression of 7 genes related to the main GO categories. (H) GSEA analysis illustrating the hallmark gene set mitotic spindle as the main factor enriched in our RNA-seq experiment. (I) Our analysis identified biological processes affected by the EFF-primed cells in the somatosensory cortex and highlighted the gene Peroxisome-Proliferator Activated Receptor alpha *(PPAR* ) as a potential target molecule underlying the therapeutic effect.

### EFF-primed fibroblasts induce metabolic signatures consistent with enhanced vascular support in the AD cortex

To identify biological processes underlying the response to EFF-primed fibroblasts, we analyzed the transcriptome of the somatosensory cortex across all four experimental groups using RNA-seq. PCA revealed clear segregation by genotype (WT vs. 3xTg-AD), with more modest variation associated with treatment (Sham vs. EFF) **(Supp. Fig. 7A).** In WT mice, *EFF* treatment resulted in 224 downregulated and 52 upregulated genes relative to sham controls **(Fig. 6A; Supp. Fig. 7B; Supp. Table 7). GO** analysis identified alterations in glucose metabolism, fatty acid oxidation, and nucleoside phosphate signaling **(Fig. 6B,C),** although these categories were driven by a small subset of DEGs. Consistent with this, GSEA Hallmarks revealed enrichment of metabolic pathways, including *Fatty Acid Metabolism, Glycolysis,* and *Oxidatiυe Phosphorylation,* indicating that healthy cortex responds to EFF-primed cells through modulation of metabolic gene programs **(Fig. 6D; Supp. Table 8).**

**Figure 7.**
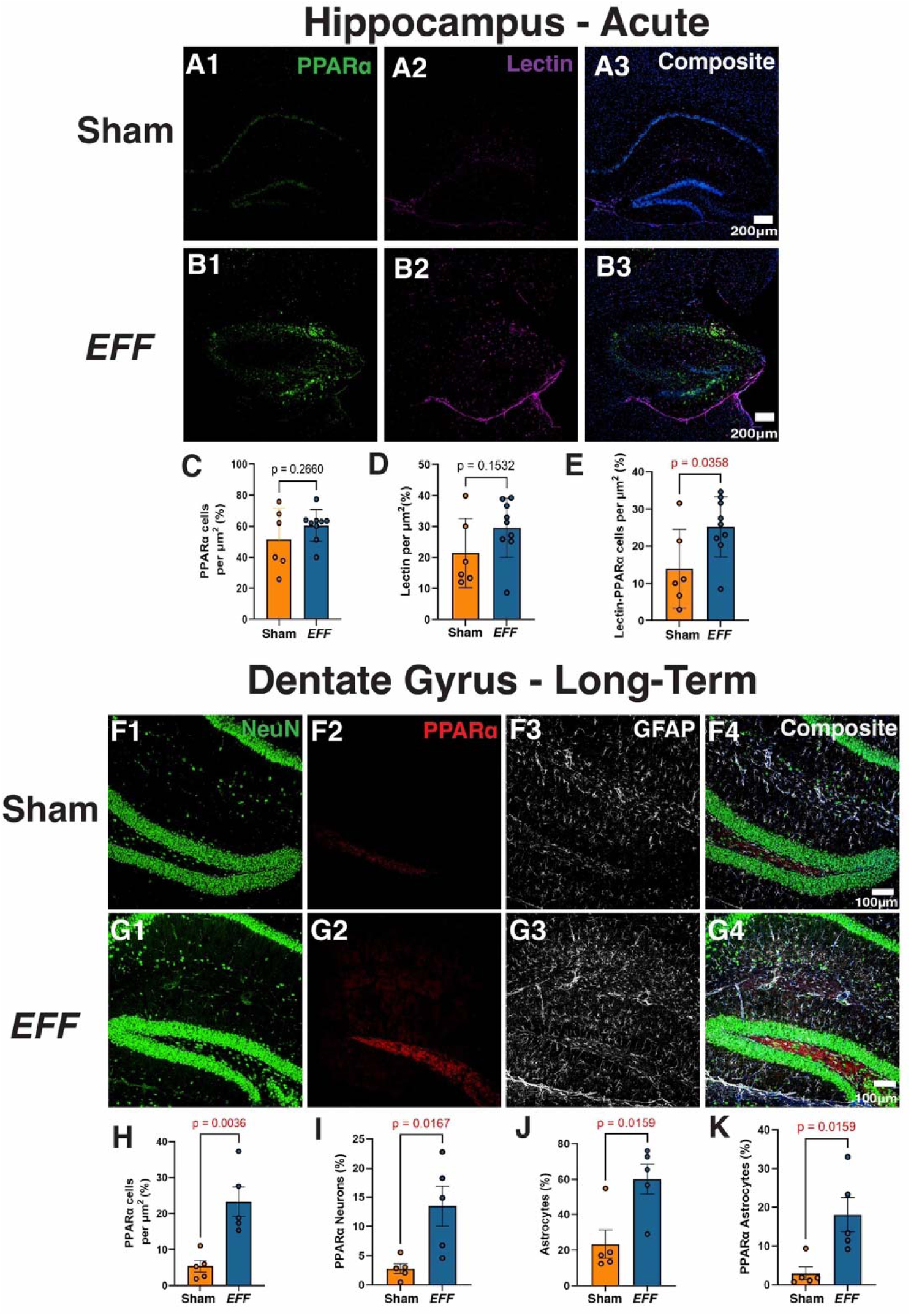
Single and multiple intracranial injections of EFF-primed cells induce an increased expression of PPARØ in the hippocampus of 3xTg-AD mice. Immunofluorescence staining for PPARD (green), lectin (magenta), and DAPI (blue) in the hippocampus of 3xTg-AD mice 7 days after a single injection with sham- **(A)** or *EFF-* primed cells **(B).** Quantification of **(C)** the number of PPARD positive cells per µm^2^, **(D)** lectin positive cells per µm^2^, and **(E)** PPARD and lectin double positive cells per µm^2^. Immunofluorescence staining for NeuN (green) PPARD (red), GFAP (white), and DAPI (blue) in 3xTg-AD mice 6 weeks after the last injection of the 3 scheduled monthly injections with sham- **(F)** or EFF-primed cells **(G).** Quantification of **(H),** the number of PPARD positive cells per µm^2^, **(I)** percentage of NeuN and PPARD double positive cells, **(J)** GFAP per µm^2^, and **(K)** percentage of GFAP and PPARD double positive cells (n= 5-8. Student’s t-test).

In 3xTg-AD mice, the transcriptional response to *EFF* treatment differed markedly. Here, we identified 76 upregulated and 4 downregulated genes relative to sham-treated AD mice **(Fig. 6E; Supp. Fig. 7C; Supp. Table 9).** GO terms again highlighted pathways related to fatty acid oxidation, glucose and carbohydrate metabolism, and ribonucleotide biosynthesis **(Fig. 6F,G),** driven largely by upregulation of genes such as *Msti, Irsi, Ppara, Uprt, mt-Atp8, Gapdh,* and *Ppargc1b.* Notably, GSEA Hallmark analysis identified the *Mitotic Spindle* as the only significantly enriched process **(Fig. 6H. Supp. Table 10),** suggesting a more discrete transcriptional signature in the AD brain. Given the reduction in CBF observed in 3xTg-AD mice and the increased vascular area induced by EFF-primed fibroblasts, these metabolic gene changes likely reflect an EFF-driven restoration of vascular support and energy delivery. Additional pairwise comparisons between genotypes and treatment groups **(Supp. Fig. 7D-I)** further highlighted differences in inflammatory, metabolic, and angiogenic pathways, with *EFF* treatment in AD mice uniquely enriching processes linked to inflammation, angiogenesis, and cell cycle regulation.

In summary, transcriptomic profiling of the somatosensory cortex reveals that EFF-primed fibroblasts induce metabolic gene programs, particularly those related to fatty acid oxidation and glucose metabolism, in both WT and 3xTg-AD mice. In the AD brain, where vascular dysfunction and impaired energy homeostasis are prominent, these changes align with the observed increase in cortical vascularization and may represent a mechanism by which *EFF* treatment improves vascular and cognitive function.

### EFF-primed fibroblast therapy induces cell-type—specific upregulation of Ppara in vascular and hippocampal compartments

Closer examination of the GO analyses for both WT and 3xTg-AD mice revealed shared transcriptional responses to EFF-primed fibroblast therapy, including upregulation of *Irsi, Uprt, Ppargcib,* and *Ppara.* Because glucose metabolism and fatty acid oxidation emerged as the principal cellular processes altered by *EFF* treatment, we next sought to identify molecular regulators that could account for the therapeutic effects observed in 3xTg-AD mice. This analysis highlighted peroxisome proliferator-activated receptor alpha *(Ppara)* as a key candidate. *Ppara* encodes a nuclear hormone receptor activated by peroxisome proliferators, which subsequently induces genes involved in fatty acid β-oxidation, insulin signaling, and related metabolic pathways, processes that enhance metabolic efficiency, support cell survival, and reduce apoptosis ^72^. Notably, previous studies have reported multiple beneficial roles for *Ppara* in AD, including inhibition of amyloidogenic processing, attenuation of tan phosphorylation, suppression of neuroinflammation and oxidative stress, activation of autophagy, and modulation of nerrrotransmission73 **(Fig 61) .**

To further characterize Ppara expression in response to EFF-primed fibroblast therapy and determine whether vascular cells contribute to these changes, we examined PPARα protein levels and lectin by immunostaining in the hippocampus of 3xTg-AD mice harvested 7 days after injection (acute treatment) **(Fig. 7A,B).** Although the total number of PPARα^+^ cells and lectin^+^ vascular cells did not differ between sham- and EFF-treated mice **(Fig. 7C,D),** we observed a significant increase in PPARα^+^/lectin^+^ double-positive cells in the EFF-treated group **(Fig. 7E),** indicating enhanced PPARα expression specifically within vascular-associated cells at this early time point.

Analysis of long-term treated mice (three EFF-primed cell injections; **Fig. 7F,G)** in the dentate gyrus revealed broader cellular changes. *EFF* treatment increased the total number of PPARα^+^ cells **(Fig. 7H),** as well as PPARα expression in neurons (NeuN^+^; **Fig. 7I)** and astrocytes (GFAP^+^; **Fig. 7J,K)**, suggesting a more widespread upregulation of PPARα in hippocampal circuits over time.

In the somatosensory cortex, acute analyses showed an increase in lectin^+^ vascular cells **(Supp. Fig. 8A-C)**, consistent with early pro-vascular effects of *EFF* treatment. However, neither the total number of PPARα^+^ cells nor the number of PPARα^+^ vascular cells differed between treatment groups at this stage **(Supp. Fig. 8D,E).** In long-term treated mice, western blot analysis of cortical lysates revealed no overall change in PPARα protein levels **(Supp. Fig. 8F,G)**, and complementary IHC analyses showed no differences in PPARα^+^ neurons or astrocytes **(Supp. Fig. 8H-K).**

Taken together, these results indicate that EFF-primed fibroblasts induce localized and cell-type-specific increases in PPRα expression, with early upregulation in vascular-associated cells and sustained effects in select hippocampal neuronal and glial populations. These transcriptional and cellular changes align with the metabolic pathways identified in our RNA-seq analysis and may contribute to the therapeutic benefits observed in 3xTg-AD mice.

## Discussion

In this study, we demonstrate for the first time that pro-vasculogenic fibroblasts generated through direct nuclear reprogramming hold therapeutic potential for AD. AD is characterized by the convergence of multiple pathological processes, obscuring any singular etiological driver^74^. While the amyloid cascade hypothesis has long dominated mechanistic thinking^75–77^, proposing that aberrant Aβ processing initiates downstream neurodegeneration^78^,^79^, its role has been increasingly challenged. Despite associations between Aβ, cerebral amyloid angiopathy(CAA)^80^, BBB dysfunction, and reduced CBF^7–81^, decades of Aβ-centric interventions have yielded limited clinical benefit ^82^. Only recently have anti-Aβ antibodies such as Donanemab and Lecanemab shown modest improvements^83^, yet Aβ burden itself correlates poorly with cognitive decline^84–88^, underscoring the need for alternative therapeutic strategies.

Although additional hypotheses such as tan hyperphosphorylation, calcium dyshomeostasis, cholinergic dysfunction, etc., offer partial explanations^89^, none fully account for the early and systemic features of AD. In contrast, the vascular hypothesis suggests that cerebrovascular dysfunction precedes and drives subsequent amyloid and tan pathology, neurodegeneration, and cognitive impairment^7–81–90–92^. This is supported by multiple lines of evidence, including early CBF reductions and BBB breakdown in patients, postmortem findings of reduced vascular density, and experimental studies showing that vascular injury impairs Aβ clearance^18^, accelerates AD-related pathology^19^, and produces AD-like cognitive decline^93^. The two-hit vascular model further proposes that early vascular/BBB disruption initiates a cascade culminating in the accumulation of plaques and tangles^24,94^. Given this, the cerebrovascular compartment has emerged as a compelling therapeutic target^95–97^, suggesting that interventions capable of restoring cerebrovascular function may modify the course of AD more effectively than approaches targeting downstream protein aggregates. Our results place vascular-focused, cell-based therapy within this emerging therapeutic landscape, supporting the concept that augmenting cerebrovascular health can meaningfully influence AD-related outcomes.

To address the cerebrovascular contribution to AD, we implemented a non-viral direct reprogramming strategy to generate vasculogenic fibroblasts capable of counteracting the marked reductions in vascular function characteristic of the disease^98^. This approach relies on electrotransfection-mediated overexpression of lineage-defining transcription factors, enabling rapid redirection of somatic cell identity. Our group has pioneered non-viral vasculogenic reprogramming for a wade variety of applications^50–99–101^ Using this method, we recently demonstrated that the *Etυ2, Foxc2,* and *Fli1 (EFF)* transcription factor cocktail efficiently primes mouse and human fibroblasts toward endothelial-like phenotypes (iECs), enabling therapeutic vascularization in models of ischemic skin^50–101^, peripheral nerve injury^100^, and stroke^51^

Here, we evaluated this vasculogenic cell therapy to the extensively characterized 3xTg-AD model, which recapitulates key AD pathologies, including early CBF decline. Importantly, while previous work mainly focused on fully reprogrammed iECs, the early post-transfection period remained unexplored. Our RNA-seq analysis revealed that by 24 hours after *EFF* delivery, fibroblasts already upregulate genes involved in vasculogenesis, angiogenesis, and endothelial adhesion **(Fig. 1)**, indicating rapid commitment to a vascular program. Concurrently, EFF-transfected cells released EVs enriched in *EFF* factors capable of propagating reprogramming signals to neighboring cells **(Supp. Fig. 2).** These findings motivated us to deliver EFF- primed fibroblasts shortly after transfection to leverage both early intrinsic transcriptional remodeling and EV-mediated paracrine amplification. Our *in υiυo* studies show that a single ventricular injection of EFF-primed fibroblasts in 7-m0nth-0ld 3xTg-AD mice increases CBF within one week **(Fig. 2B,C)**, consistent with prior observations in wild-type mice^51^ EFF-primed cells also improved neurovascular coupling in the barrel cortex **(Fig. 2D-G)**, suggesting restoration of NVU function likely driven by direct vasculogenic activity and paracrine EV transfer. Notably, whisker-evoked responses were asymmetrical, a finding aligned with reports of hemispheric variability in cerebrovascular reactivity in AD and potentially reflecting regional heterogeneity in treatment distribution.

Because 3xTg-AD mice exhibit cognitive impairment, gliosis, synaptic deficits, and early Aβ deposition by 6-7 months ^68^,^102^, we next evaluated whether early and repeated EFF-primed cell delivery (initiated at 4 months) could modify cognitive trajectories typical of AD^103–105^. Serial injections preserved spatial learning and memory in 3xTg-AD mice but did not alter cognition in WT animals **(Fig. 3)**, strongly supporting the vascular hypothesis of AD and demonstrating that targeting the vascular compartment can modify disease-relevant behavior. Recognition memory remained unaffected; however, 3xTg-AD mice at this age exhibit minimal NOL/NOR deficits^106^, aligning with our observations. Lastly, although synaptic plasticity deficits are well documented in 3xTg-AD mice^68–70^, *EFF* therapy did not rescue hippocampal LTP **(Supp. Fig. 4).** However, LTP does not always correlate with behavioral memory performance^107^,^108^, and EFF-induced cognitive improvement may instead reflect enhanced circuit-level function, engram stability, or extra-hippocampal contributions.

We delivered cells into the lateral ventricles to maximize distribution via CSF flow and perivascular pathways^109^. BrdU^+^ cells were detected in cortical and hippocampal regions distal to the injection site, including the ependyma, somatosensory cortex, and dorsal hippocampus **(Fig. 4)**, indicating long-range dispersal and survival for at least six weeks. Co-localization with lectin demonstrated close association with host vasculature across regions. EFF-primed cells increased vascular area selectively in 3xTg-AD mice **(Fig. 4H)**, but not in WT mice, consistent with a disease-context-dependent response. Together, these findings indicate that EFF-primed fibroblasts produce a transient increase in CBF and a more durable expansion of vascular architecture in AD brains. Key differences between disease, non-disease context may be in part explained by different metabolic needs that lead to the increase of key factors controlling insulin signaling and lipid oxidation such as *Pparα*.

Aβ pathology also responded to therapy. In the somatosensory cortex, EFF-treated mice exhibited reduced Aβ load within layers IV-V **(Fig. 5).** Given the dynamic relationship between intracellular and extracellular Aβ^110^ and reports that restoring CBF via reduced neutrophil adhesion lowers Aβū-ūū independently of plaque burdeni, our findings suggest improved Aβ clearance rather than direct inhibition of production. Enhanced CBF and expansion of vascular area, among other things, may contribute to this effect^111^. Corresponding analyses of astrogliosis and microglial activation (GFAP, IBA1, CD68) revealed no treatment-induced differences at the ages examined **(Supp. Fig. 6).** Future studies at later disease stages (12-16 months) will clarify whether vascular repair influences neuroinflammation.

Transcriptomic profiling of somatosensoiy cortex revealed that *EFF* therapy regulates glucose and lipid metabolism in both WT and 3xTg-AD mice **(Fig. 6).** Interestingly, these pathways were driven by a small number of DEGs, with the majority of differentially expressed transcripts lacking well-characterized functions. Strikingly, genes involved in insulin signaling, glucose uptake, and mitochondrial metabolism exhibited opposite regulation in AD vs. WT brains, with the *EFF* treatment upregulating these pathways in 3xTg-AD mice but downregulating them in WT mice. These divergent responses likely reflect distinct energetic states, such as homeostatic neurovascular coupling in WT brains versus impaired glucose utilization and mitochondrial function in AD. In AD mice, improved cerebrovascular function may restore metabolic support, prompting adaptive upregulation of energy-related genes.

Among the metabolic regulators identified, PPARα emerged as a key candidate. PPARα governs lipid β-oxidation, insulin signaling, and mitochondrial efficiency, and has been implicated in suppressing Aβ synthesis, tan phosphorylation, neuroinflammation, and oxidative stress^73^. Our immunostaining revealed increased PPARα protein in specific hippocampal populations, reinforcing its potential role in mediating therapeutic benefit. Indeed, clinical testing of the PPARα agonist Gemfibrozil in prodromal AD demonstrated favorable safety and early signals of efficacy^112^, and other agonists (e.g., fenofibrate, WY-14643, GW7647) continue to be investigated^113^,^114^. Future studies combining selective modulation of PPARα with EFF-primed fibroblasts will be essential to determine whether this pathway represents a mechanistic driver of the observed vascular and cognitive improvements.

Critically, the therapeutic translation of these pre-clinical findings is readily achievable. Administration of EFF- primed fibroblasts into the ventricular system can be accomplished using well-established, minimally invasive neurosurgical techniques^115^. Specifically, stereotactic placement of an infusion cannula into the lateral ventricle allows for precise deliveiy of this cell therapy. This approach mirrors existing infusion paradigms currently employed for drug administration to the intraventricular system. Acute or repeated infusions can be performed through chronically implanted subcutaneous injection reservoirs, (e.g., Ommaya reservoir), which provide reliable access for ongoing therapeutic interventions^116^.

## Conclusions

Our findings reveal that fibroblasts primed with *EFF* rapidly transition toward a vascular phenotype, exhibiting angiogenic transcriptional signatures and releasing EVs that cany the same instructive factors. These EVs provide a plausible mechanism for amplifying and disseminating the reprogramming signal both *ex υiυo* and within the brain microenvironment. *In υiυo* delivery of EFF-primed cells produced acute improvements in CBF and neurovascular coupling, hallmarks of restored vascular responsiveness and metabolic support. These functional gains were accompanied by reduced spatial memory deficits, increased vascular area, and long-range dissemination of the injected cells, with histological evidence suggesting potential vascular integration. The treatment also reduced intracellular Aβ burden, an effect consistent with diminished early neurotoxicity and improved proteostatic handling. Transcriptomic analyses further revealed coordinated shifts in metabolic gene expression in both WT and AD mice, converging on pathways associated with brain energy expenditure. Notably, elevated *Pparũ* expression at the mRNA and protein levels identifies this factor as a candidate mediator linking vascular commitment, metabolic adaptation, and functional recovery. Together, these results demonstrate that *EFF* priming initiates a rapidly propagating vascular program capable of reshaping neurovascular function, cellular homeostasis, and molecular signaling in the AD brain.

## Methods

### Experimental design

The main goal of this study was to examine whether the intracranial injection of EFF-transfected fibroblasts into a mouse model of Alzheimer’s Disease could enhance cerebral blood flow, increase vascularity, and reduce disease burden. It is considered an exploratoiy and hypothesis-generating study. No power analysis was performed to predetermine the sample size. For *in vitro* and *in vivo* studies, biological replicates and end points were determined based on previous reports. All data collection, processing, and analysis were performed by investigators blinded to the sample experimental group. In addition, for *in υiυo* studies animals for each of the 2 genotypes were randomly assigned within each cage to a single treatment group (i.e., sham- or *EFF-* transfected cells), securing this way equal representation for each genotype and treatment. Successful survival to intracranial injection throughout the experiment was a prerequisite for inclusion into the experimental analysis. Post hoc exclusion criteria included motivation to perform the behavioral task, photomicrographs with observed artifacts and/or statistical outliers (Tukey Method).

### Animal husbandry

The mouse strain used for this research project, B6;129 Tg(APPSwe,tauP301L)1Lfa *Psen1^tm1Mpm^*/Mmjax, RRID:MMRRC_034830-JAX, (3xTg-AD)^68^,^117^,^119^ was obtained from the Mutant Mouse Resource and Research Center (MMRRC) at The Jackson Laboratoiy, an NIH-funded strain repositoiy, and was donated to the MMRRC by Frank Laferla, Ph.D., University of California, Irvine. Mark P. Mattson, Ph.D., Johns Hopkins University, School of Medicine. As controls we used the WT mouse strain B6129SF2/J_ioio45 RRID:IMSR_JAX:loio45 also obtained from The Jackson Laboratory. Eight-week-old male and female mice from both strains 3xTg-AD and B6129SF2/J controls were purchased to initiate breeding colonies. From these colonies animals from litter born in house were allocated for the experimental procedures. All animals were maintained on a 12h light: dark cycle with constant humidity and temperature, and with ad libitum access to food and water. All animal experiments were performed in accordance with protocols approved by the Laboratoiy Animal Care and Use Committee at The Ohio State University (IACUC no. 2016A00000074-R2 and IACUC no. 2009A0006-R5).

### Cell electrotransfection

To induce vasculogenic cell reprogramming, primaiy Mouse Embryonic Fibroblasts (pMEFS) (Millipore-Sigma, Cat. PMEF-HL) were transfected with expression plasmids encoding 3 transcription factors *(Etυ2, Foxc2, Fli1)* each fused to the reporter Green Fluorescent Protein (GFP) and under the control of a cytomegalovirus (CMV) promoter. As control we used an expression plasmid encoding only the GFP reporter (Sham plasmid) also under control of the CMV promoter. A full list of plasmids used can be found in Supp. Table 11. All plasmids were used to transform DHsū competent bacteria (Invitrogen. Cat. 18265017) using the heat shock protocol. Briefly, 50µL of DHsū cells were thawed on ice, combined with 4µL of long/µL plasmid DNA and incubated for 20 minutes on ice. Heat shock was set by incubating the cells in a water bath at 42 □ for 2 minutes and immediately transfer to the ice bath. Two minutes after the shock SOC media was added to the cells and after incubation at 37□, cells were plated in LB-Agar prepared with Ampicillin as selection antibiotic. After selection of colonies of transformed bacteria, plasmids were prepared using a mid- capacity plasmid DNA isolation kit (ZymoPURE, Cat. D4201) following the manufacturer’s instructions. Plasmid DNA concentrations were obtained with a NanoDrop™ 2000/2000c Spectrophotometer (Thermo Fisher Scientific, Cat. ND-2000). For specific tracking of cells after specified intracranial injection, cells were prelabeled with 5-Bromo-2’- Deoxyuridine (BrdU) (Sigma, B5002-1G) by exposing cell cultures in logarithmic growing phase to íouM BrdU for 48h just before electroporation. For electroporation we used the Neon™ transfection system (Thermo Fisher Scientific MPK5000). For this, plasmids were prepared at concentrating of 50ng/µL in Neon buffer R™. The *Etυ2, Foxc2, Fli1 (EFF)* cocktail was mixed at 1:1:1, whereas the sham plasmid was prepared at 150ng/µL to match the EFF cocktail mass. Primaiy Mouse Embiyonic Fibroblast were grown in Dulbecco’s Modified Eagle’s Medium (DMEM) (Corning, Cat. 10-013-CV) supplemented with 10% exosome-depleted fetal bovine serum (FBS) (Thermo Fisher Scientific, Cat. A2720801). Before electroporation pMEFs were harvested and resuspended in the mixture of plasmid plus buffer R™. After resuspension cells were electroporated with a single pulse of 1425mV and 30ms of duration. Then electroporated cells were incubated for 24h before harvesting for injection.

### Extracellular Vesicles uptake

To probe if after cell electroporation released EVs are loaded with our 3 reprogramming factors, cells were electroporated and plated with DMEM media supplemented with 10% exosome-depleted FBS on top of a 12-multiwell plate insert fabricated with sterile PET mesh with a pore size of 0.4 µm. This pore size allowed the free exchange of EVs between the insert and the bottom chamber of the multiwell plate but prevents the migration of cells. In the bottom chamber naive, non-transfected cells were plated simultaneously with DMEM media supplemented with 10% exosome-depleted FBS **(Fig. 1).** Twenty-four hours after plating, cells, and conditioned media EVs isolated with Total Exosome Isolation Reagent (Invitrogen, Cat. 4478359) from inserts and bottom chambers were processed for RNA purification.

### RNA isolation and qRT-PCR

Cells and EVs were processed for total RNA purification using TRIzol reagent (Thermo Fisher Scientific, Cat. 15596026) following the manufacturer’s protocol. After purification, RNA samples were treated with DNAse I to remove any trace of contaminant DNA. Total RNA was quantified NanoDrop™ 2000/2000c Spectrophotometer (Thermo Fisher Scientific, Cat. ND-2000). Following RNA quantification cDNA from each sample was prepared by reverse transcription using SuperScript IV VILO Master Mix (Thermo Fisher Scientific, Cat. 11756500), making sure that the final concentration of cDNA was equal for all the samples. Alternatively, RNA from transfected cells or brain tissue was processed for RNA-sequencing using TRIzol reagent for isolation followed by a column-based kit for purification and cleaning (Thermo Fisher Scientific. Cat. 12183018A). For quantitative RT-PCR TaqMan primers (Thermo Fisher Scientific) were employed and reactions were run and quantified in a QuantStudio 3 machine (Thermo Fisher Scientific). TaqMan primers utilized in this study are detailed in Supp. Table 12. Data analysis was carried out following the method outlined by Königshoff et al^120^, where the ΔΔCt values represent the logarithm of the fold change.

### Intracranial cell Injection

All the intracranial injections were performed at The Preclinical Therapeutics Mouse Modeling Shared Resource (PTMMSR) from the Comprehensive Cancer Center (CCC) at The Ohio State University Medical Center. Twenty-four hours after electroporation with the *EFF* plasmid cocktail or the sham plasmid, cells were intracranially injected via stereotaxic surgery. Surgeries were performed at 4-, 5- and 6-months of age targeting both LV [coordinates from bregma: antero-posterior (AP): -0.32 mm; medio-lateral (ML) ± 1.0 mm; dorso- ventral (DV) -2.0 mm]. On the day of surgery, mice were deeply anesthetized with isoflurane (induced at 5%; maintained at 1.5 to 2%) confirmed by toe pinch test before the start of any procedure. Mouse head was shaved and secured to stereotactic frame. After thorough cleaning of the head skin with povidone-iodine solution and alcohol, skull was exposed and motorized drill was used for trepanation at the indicated coordinates in both hemispheres. Using 2 inches and 32-gauge needle with a 10-µl Hamilton syringe connected to a nano-injector, electroporated cells were delivered at the specified DV coordinate in a suspension 8 µl of standard DMEM at a concentration of 5xio^4^ cells µL and a rate of 2 µl/min. After the entire volume of cell suspension was injected, the needle was left in place for additional five minutes to avoid backflow of injectate. After the procedure was finished, the skin was closed again using a sterile nylon suture reinforced with tissue-compatible glue. For recovery mice received toouL of pre-warmed sterile saline solution and were monitored for 3h.

### Behavioral analysis

All behavioral assessments for this study were performed at the Rodent Behavior Core from the Department of Neuroscience at The Ohio State University Medical Center. Two weeks after the last stereotaxic injection, recognition and spatial memory were evaluated by Novel Location/Recognition task (NOL/NOR) and Barnes Maze, respectively. For the NOL/NOR task, mice were placed in an empty testing chamber (45 x 24x 22 cm rodent cage with bedding) for 10 minutes on day 1 of habituation. On testing day, day 2, 3 sessions occur about one hour apart. For session 1, mice were placed in the cage with 2 identical objects for 10 minutes. Mice were then removed and returned to their home cage. After a 1-h0rrr interval, mice were returned to the testing cage for 10 minutes, which contains the same 2 identical objects with one moved to a novel location. One hour later, for session 3, the mice were placed back into the testing cage which now contains one of the original objects and a novel object. All 3 sessions are recorded by the ANYmaze tracking system. Time interacting with each object was scored manually following testing. The chamber was cleaned with 70% ethanol between mice. For sessions 2 and 3, recognition index was calculated for each mouse following the formula RI = (T_ne_w/Tt_o_tai)*ιoo, where T_new_ = time exploring the new location/object and T_to_tai= total exploration time. Barnes maze is a memory task that requires mice to use spatial cues around an elevated platform to locate a hidden goal box among multiple holes located around the platform. The animals were exposed to bright light (clamp lamp positioned above the maze, 75W) throughout testing and white noise (66 dB) while on the maze. To help increase the motivation to enter the escape box, bedding from each cage was placed inside. The escape box was cleaned and replaced with the next cage’s bedding in between. Each mouse was placed in the middle of the maze at the start of each trial and allowed to explore for 2 minutes. Each trial ended when the mouse entered the escape box or after 2 minutes had elapsed. Immediately after the mouse entered the box, it was allowed to stay there for about 30 seconds. If the mouse did not reach the goal within 2 minutes, the experimenter gently guided the mouse to the escape box and left the mouse inside for about 30 seconds. Once the mouse was placed back in its home cage, the maze and escape box were cleaned with 70% ethanol followed by testing of the next mouse. This was repeated until each animal had received 3 trials per day over 5 consecutive days. During these training days, the latency, distance traveled, and time in goal perimeter were recorded by the ANY-maze tracking system (Stoelting). The number of errors, counted manually eveiy time the animal’s nose fully entered the wrong hole, were determined after the trial ended. On day 6, the probe trial was conducted. The acquisition index was determined by subtracting the latency time on the 3^rd^ trial of each day from the latency time on the 1^st^ trial on the same day. Similarly, the saving index was calculated by subtracting the latency on the 1^st^ trial of the day from the latency on the 3^rd^ trial from the day before. During the probe trial, the escape box was removed, and the mouse was placed in the middle of the maze and allowed to explore for a fixed interval of 60 seconds. The same parameters plus distance and time spent in the goal quadrant were quantified automatically by the ANY-maze tracking system (Stoelting), number of errors was quantified manually.

### Laser speckle imaging

Brain perfusion in WT and 3xTg-AD mice was assessed 7 days after cell intracranial injections. After mice were deeply anesthetized with isoflurane vaporization induced at 5% and maintained at 2%, head skin was cleaned with povidone-iodine solution and alcohol and a longitudinal incision was made to expose the skull. Sterile saline solution was used to keep the skull moisturized and clean of debris and hairs. Laser speckle imaging (LSI) recordings were taken using the moorO2Flo perfusion and oxygenation imager (Moor instruments) with a sampling rate of 20 frames per second and a resolution of loµm/pixel. During the session, the exposed skull was 5-8 cm below the imager. The first minute of recording was used to allow the stabilization of respiratory and cardiac frequencies. Perfusion recordings were then taken for 20s and flux (A.U) was reported as the average perfusion levels in these last 20s. For neurovascular coupling (NVC) assessment, mice preparation and recording settings are the same as described above. After one minute of stabilization, recordings were taken under the following protocol: one minute of baseline perfusion, 30s of recording while mechanically stimulating the right whiskers with a paintbrush, baseline for one minute, 30s of recording while stimulating the left whiskers, baseline for one minute, 30s of recording while stimulating the right whiskers, baseline for one minute, and 30s of recording while stimulating the left whiskers.

### Immunostaining

After harvesting, brains were transferred to ice-cold HBSS (StemCell Technology, Cat. 37150) and separated in two with a single cut along the midline. From one hemisphere, the hippocampus and somatosensoiy cortex were flash-freeze in diy ice and stored at -80 □ for protein isolation and biochemical analysis. The second hemisphere was fixed in cold 10% formalin (Sigma, Cat. HT501-128) for 48h, followed by incubation with sucrose 30% prepared in PBS for ciyopreservation and finally embedded in Optimum Cutting Temperature O.C.T compound (Fisher Scientific, Cat. 4585) for ciyo-sectioning. For immunofluorescence analysis 40µm free floating sections were obtained with a ciyostat and preserved in 0.01% sodium azide in PBS. To be able to compare similar tissue sections from each mouse, brains were ciyo-sectioned in a series of 6 sister wells. For BrdU staining antigen retrieval was achieved by incubating the free-floating sections in 2N HC1 for ih at 37°C and washed with boric acid 50mM pH 8.0 for 20 minutes at room temperature. Sections from each mouse were washed and permeabilized in 1% Triton X-100 (Ricca Chemicals, Cat. 8698.5-11.6) in PBS, followed by blocking with 10% Normal Goat Serum NGS (Vector Labs, S-1000) in PBS. Primaiy antibodies were incubated in blocking solution for 16h at 4°C, followed by washes and incubation for 2h with secondary antibodies conjugated to a fluorophore prepared in 5% NGS. When required Lycopersicon Esculentum Lectin (LEL, TL) conjugated to DyLight® 48 at 1:20 was used label blood vessels (Vector Labs, Cat. DL-1174-1). This was followed by incubation with 4’,6-diamidino-2-phenylindole (DAPI) (Invitrogen, Cat. D3571) and mounted in charged microscope slide using mounting media (Vector Labs, Cat. H-1700). Imaging started 24h after mounting. For immunohistochemical analysis free floating sections prepared as described before were treated with 88% formic acid for 20 minutes for Aβ antigen retrieval. After washing the formic acid, sections were permeabilized with 1% Triton X-100 and incubated with 10% Hydrogen peroxide (Fisher Scientific, Cat. H325- 100) prepared in methanol for 10 minutes. Blocking was done with a 10% NGS solution plus a Biotin/Streptavidin blocking solution (Vector labs, Cat. P-2001). Following blocking, sections were incubated 16h at 4 □ with the biotinylated primaiy antibody against Aβ clone 6E10 (Biolegend, Cat. 803007). Subsequently sections were incubated with an Avidin-Biotin Complex (ABC) (Vector labs, Cat. PK7100) bound to the peroxidase enzyme. Finally, all sections were exposed to the chromogen 3,3’-Diaminobenzidine (DAB) (Vector Labs, Cat. SK-4105). Sections were dehydrated in Xylenes and mounted in microscope slide with DPX mounting media (Electron Microscopy Sciences, Cat. 13512). A list of primaiy and secondaiy antibodies used in this study is detailed in Supp. Table 13. Sections processed for immunofluorescence and immunohistochemistiy were imaged using Nikon Ti2-E microscope operating on NIS-Elements AR version 5.20 and with a sCMOS camera for immunofluorescence (Hamamatsu Orca Flash 4.0.) and color digital CMOS camera for immunohistochemistiy (Nikon DS-Ri2).

### Image analysis

Quantification of immunohistochemical staining for Aβ and fluorescence signal for lectin was quantified using ImageJ Fiji software. Aβ load was quantified from color images taken from immunohistochemistiy processed sections. Images were loaded into ImageJ Fiji and converted into 8-bit grayscale images. Specific ROI areas were selected by a blind investigator including the entire somatosensory cortex in the section and excluding section artifacts such as tears and debris. Images were then transformed into binaiy images using the MaxEntropy method. From each image the total *area* of the ROI and the *percentage area.* The percentage area represents the proportion of the ROI area with positive labeling. The calculated percentage area was average from 2 sections per mouse. The same approach was used for processing of lectin fluorescence images but instead of *percentage area,* the total area positive for lectin signal was calculated and used for quantification analysis. For analysis and quantification of immunofluorescence signal for Iba1, CD68 and GFAP images were acquired on a Nikon Eclipse Ji confocal microscope controlled by the NIS-Elements AR version 6.02.01 (Nikon instruments). Images were taken using a 20x objective taking multiple 1µm z-stack images for each x-y plane and channel. Once the images were loaded in the software and signals from each filter channel were separated. A binaiy mask was applied from the fluorescent signal from each channel. Image segmentation was applied to each channel, and the segmented file was used to quantify intensity and morphological parameters. For Iba1 and CD68 co-immunolabeling images, segmented files from the separated channels were superimposed to calculate the percentage of segmented cells that are positive for both Iba1 and CD68. Microglia activation was determined by morphological changes. Using the segmented channel for Iba1, sphericity, cell surface cell area, cell volume and length of the major axis were calculated. For GFAP segmented images were used to calculate total area and signal intensity.

### RNA-sequencing

RNA-sequencing was performed and analyzed by Novogene CO, following their standard pipeline. Messenger RNA was purified from total RNA using poly-T oligo-attached magnetic beads. After fragmentation, the first strand cDNA was synthesized using random hexamer primers, followed by the second strand cDNA synthesis using dUTP for directional libraiy synthesis. This was followed by after end repair, A-tailing, ligation of adapters, digestion with USER enzyme to remove the second strand, size selection, and PCR amplification, and purification. The libraiy was checked with Qubit and real-time PCR for quantification and bioanalyzer for size distribution detection. Quantified libraries will be pooled and sequenced on Illumina platforms, according to effective libraiy concentration and data amount. Original image data file from Illumina high-throughput sequencing platforms was transformed to sequenced reads by CASAVA base recognition (Base Calling). Raw data are stored in FASTQ format files, which contain sequences of reads and corresponding base quality. Raw data was first processed through in-house Perl scripts. In this step, clean data was obtained by removing reads containing adapter, reads when uncertain nucleotides constitute more than 10 percent of either read (N > 10%), and reads when low quality nucleotides (Base Quality less than 5) constitute more than 50 percent of the read low-quality reads from raw data. At the same time, Q20, Q30 and GC content the clean data were calculated. All the downstream analyses were based on clean data with high quality. For mapping of reads to the mouse genome, the reference genome and gene model annotation files were downloaded from genome website directly. The index of the reference genome was built using Hisat2 v2.0.5 and paired-end clean reads were aligned to the reference genome using again Hisat2 v2.0.5^121^. featureCounts vl.5.o-p3^122^ was used to count the reads numbers mapped to each gene. And then FPKM of each gene was calculated based on the length of the gene and reads count mapped to this gene. Differential expression analysis of two conditions (5 biological replicates per condition) was performed using the DESeq2Rpackage (1.20.0)^123^. DESeq2 provides statistical routines for determining differential expression in digital gene expression data using a model based on the negative binomial distribution. The resulting P-values were adjusted using Benjamini and Hochberg’s approach for controlling the false discoveiy rate. Genes with an adjusted P-value <=0.05 and absolute foldchange of 2 found by DESeq2 were assigned as differentially expressed. Gene Ontology (GO) enrichment analysis of differentially expressed genes was implemented by the clusterProfiler R package^124^, in which gene length bias was corrected. GO terms with corrected P-value less than 0.05 were considered significantly enriched by differential expressed genes. Volcano plots, unbiased clustering and Venn diagrams were created in R running in the Novomagic platform. GSEA analysis was performed with the GSEA software version 4.4.0. Significant differences in mouse *Hallmark Gene Sets* were determined with an FDR q-val <0.25

### Brain slice and fEPSP recordings and Long-Term Potentiation

Mice were anesthetized with isoflurane before decapitation. The brain was rapidly removed and placed in an ice-cold cutting solution containing the following (in mM): Sucrose 250, D-Glucose 25, KC1 2.5, NaHCO3 24, NaH2PO4 1.25, CaCl2 2.0, MgSO4 1.5 (pH; 7.3-7.4). Transverse Hippocampal slices (400 µm) were prepared with a Vibratome (VT1200S, Leica). Brain slices were transferred to chambers filled with artificial cerebrospinal fluid (aCSF) bubbled with 95% 02 and 5% CO2, and containing the following reagents (in mM): NaCl 124, KC1 3, NaHCO3 24, NaH2PO4 125, CaCl2 2, MgSO4 10, and D-Glucose 10, (pH; 7.3-7.4), and allowed to recover at 37Ū for 30 min and then moved to room temperature for around 1 hour. Individual Slices were transferred to a submerged chamber held on the fixed stage of Nikon E600FN upright microscope. Slices were perfused with 2-3 ml/min oxygenated aCSF drove by gravity exchange system and maintained at 34 □ during recording. For recording extracellular field excitatory postsynaptic potential (fEPSP) from CA1 area, custom-made twisted nichrome stimulation electrode was placed at Schaffer collaterals close to CA2 area. Stimulation pulse (too µs duration, every 30 s) was generated by isolator (Iso-flex, A.M.P.I.) under computer control. Recording electrode was pulled from borosilicate glass filled with aCSF (resistance 1.5-3 Mω) and placed in the stratum radiatum of the CA1. The distance between stimulation and recording electrodes is around 300 µm. for each slice, we firstly recorded an input-output (I/O) curves generated with varying intensity from 0.0-0.5 mA (100 µS duration) in 0.05 mA increments. The stimulation intensity which evokes around 40-50% of the maximum response, without emergence of population spikes, was chosen to conduct the following experiments. Paired-pulse ratio (PPR) was determined through measuring the two peak values of evoked fEPSPs with vaiying intervals (mS) 50, 100, 150, 200, and 250. The ratio was calculated as P2/P1. Synaptic field potentials were low pass filtered at 1 kHz and digitally sampled at 50 kHz with Axopatch 200B amplifier and Digidata 1440 A interface. After 20 minutes of baseline measurement (each pulse per 30 s), four high frequency stimulation (100 Hz), separated 20 s, were delivered to induce LTP and recording lasting for 60 min. All recordings were conducted with Clampex 10.6 and 11 software. The 20-80% slope of rising phase of fEPSP was measured off-line with Clampfit 11 (Molecular Devices) and normalized to averaged baseline recording (last 5 min) Responses from all sections were averaged for each animal.

### Western blot

Total protein from cortical tissue was isolated with radioimmunoprecipitation (RIPA) lysis buffer (Thermo Fisher Scientific Cat. 89 0 900), supplemented with the complete protease inhibitor cocktail (Roche, Cat. 4 0 693 0116 Oooi,), phosphatase inhibitor cocktail 2 (Sigma-Aldrich Cat. P5726-1,), and 1 mm phenylmethylsulfonyl fluoride (PMSF). All samples were incubated in lysis buffer for 15 min at 4 °C and overnight at -20 °C; samples were subsequently vortexed for 15 min, centrifuged at 10 000 RPM, 4 °C for 30 min and supernatant was recovered. Protein concentration was determined with the BCA Protein Assay Kit (ThermoFisher Scientific, Cat 23225) 20 µg of total protein was mixed with 4X Laemmli loading buffer (Biorad, Cat 1610747). Then, samples were denaturalized by heat at 85 °C for 2 min. Protein samples were separated by electrophoresis using a 4-20% Tris-glycine gel (Biorad, Cat 4568094), and transferred onto a PVDF membrane (Biorad, Cat. 1620261). Membrane was subsequently blocked with 5% BSA at room temperature for 1 h. Antibodies were diluted in TBS-T (1X) with 2.5% BSA and incubated overnight at 4 °C. Primaiy and secondaiy antibodies used are listed in Supp. Table 13. Finally, membrane visualization was done was done using a Chemidoc Imaging system (Biorad).

### Statistical analysis

Group allocation was based on a randomized permuted block design. To maintain rigor, investigators were blinded during data collection and analysis. Summaiy statistics reported as means and standard error of the mean (SEM) together with graphical display of the data was generated in GraphPad Prism (version 10). The analysis approach allowed us to test the effect of EFF (vs sham) overall and at each time point within WT and AD mouse model. For one time point data analysis, significant main effects of factors [(Genotype: control or 3xTg-AD), (Treatment: Sham or *EFF)]* and interactions between these factors was determined by two-tailed t-tests and two-way ANOVA. For longitudinal analysis, a repeated measures ANOVA was performed. Corrections for multiple comparisons were included when needed. Statistical significance was defined as P < 0.05. The number of replicates and tests employed are detailed in figure legends.

## Supporting information

Supplementary Figures and Supplemental Tables Titles

## Acknowledgements

Completion of all the stereotaxic surgeries was possible thanks to the expertise and support of The Preclinical Therapeutics Mouse Modeling Shared Resource (PTMMSR) at The Ohio State University Comprehensive Cancer Center.

