## Supplementary Figures and Supplemental Tables Titles for "Non-viral vasculogenic reprogramming restores cognition and mitigates pathology in Alzheimer’s disease"

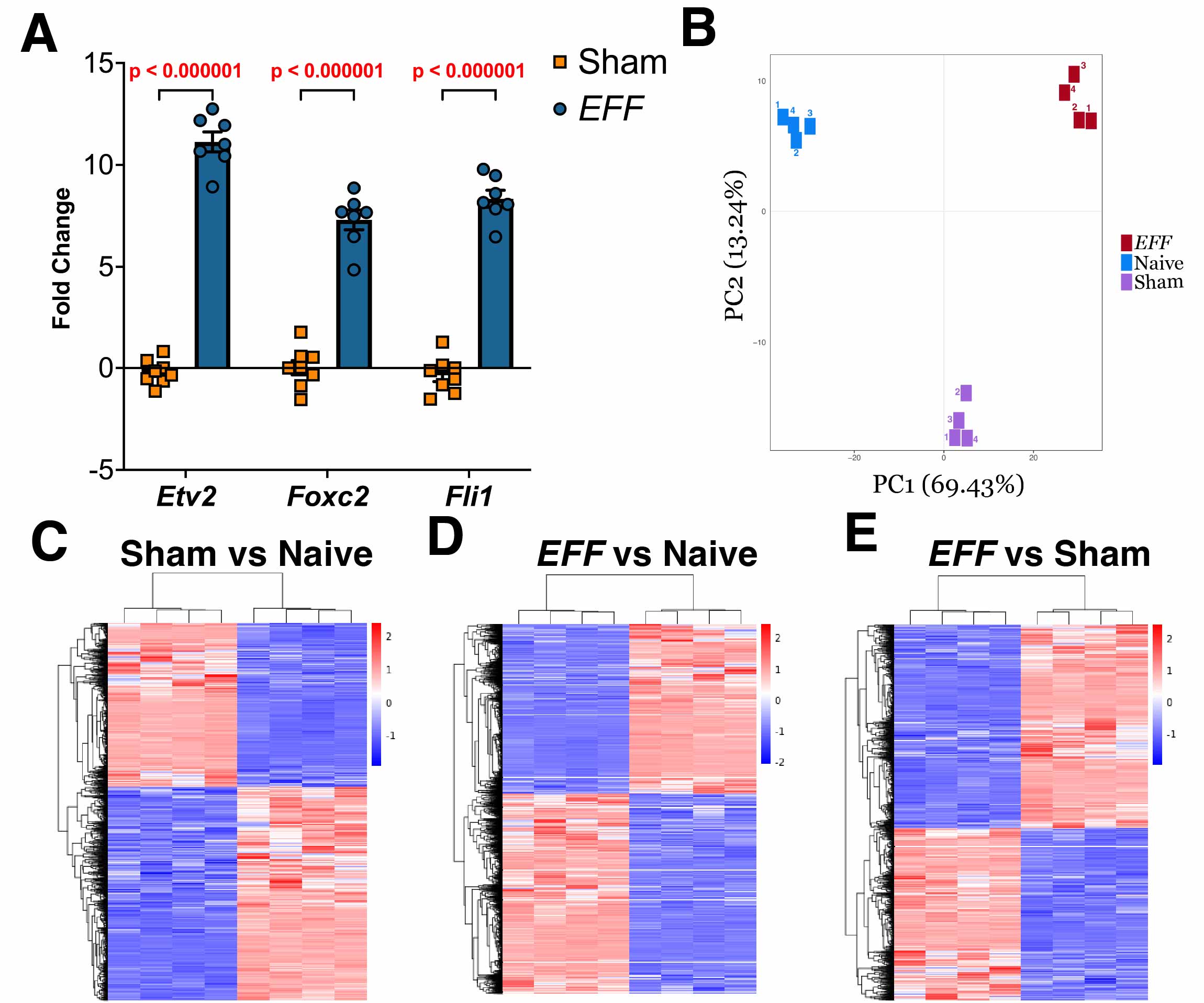

**Supplementary Figure 1. Electrotransfection of fibroblasts with the transcription factors *Etv2, Foxc2*, and *Fli1 (EFF)* induces an early vasculogenic priming. (A)** qRT-PCR analysis shows the increased expression of each reprogramming factor *Etv2, Foxc2* and *Fli1* 24h after electroporation (n=7. One-way ANOVA). **(B)** PCA plot analysis of naïve, sham plasmid electrotransfection, and *EFF-*primed cells show a clear separation between sample groups. Unsupervised heatmap clustering of DEGs between naïve fibroblasts and those electrotransfected with a sham plasmid**(C)**, naive and *EFF*-primed fibroblasts **(D),** and between fibroblasts electrotransfected with a sham plasmid and *EFF*-primed fibroblasts **(E).**

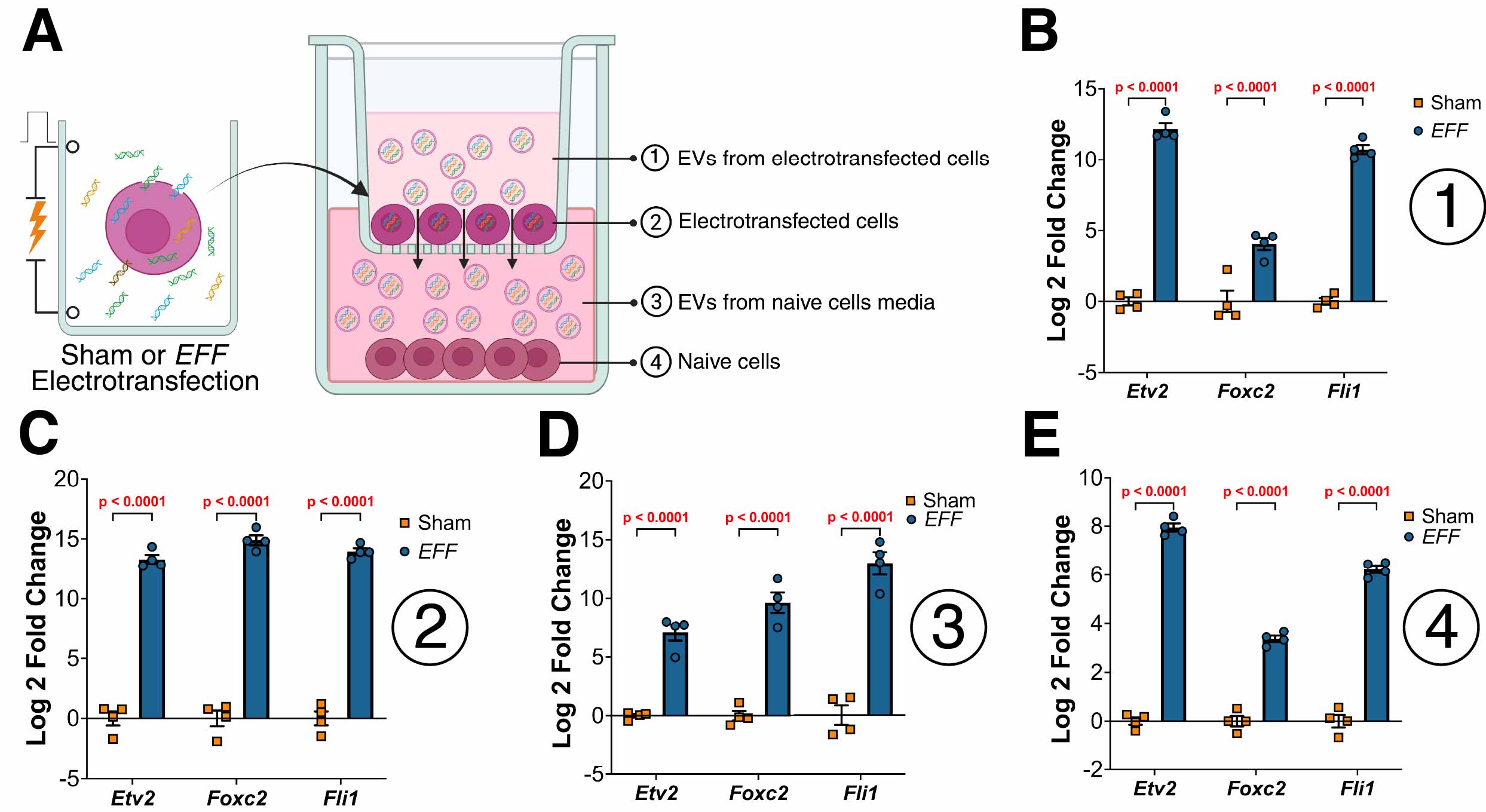

**Supplementary Figure 2. Electrotransfection of fibroblasts with the transcription factors Etv2, Foxc2, and Fli1 (EFF) induces the release of EVs loaded with the *EFF* factors. (A)** Illustration describing the EVs uptake experiment in which fibroblasts were electrotransfected with sham or *EFF* plasmids and plated on a PET insert with a pore of 0.4uM, whereas naive fibroblasts were plated in the bottom of a multiwell plate. The system was incubated for 24h and expression/presence of the reprogramming factors mRNA was probed on cells from insert and bottom chamber as well as in EVs isolated from the enriched media obtained from the insert and bottom chambers. qRT-PCR s shows presence of all the reprogramming factors in EVs purified from the **(B)** insert media, **(C)** expressed by the transfected cells, **(D)** its presence in the EVs purified from the bottom chamber media, and **(E)** expression in naïve (not electrotransfected) cells in the bottom chamber.

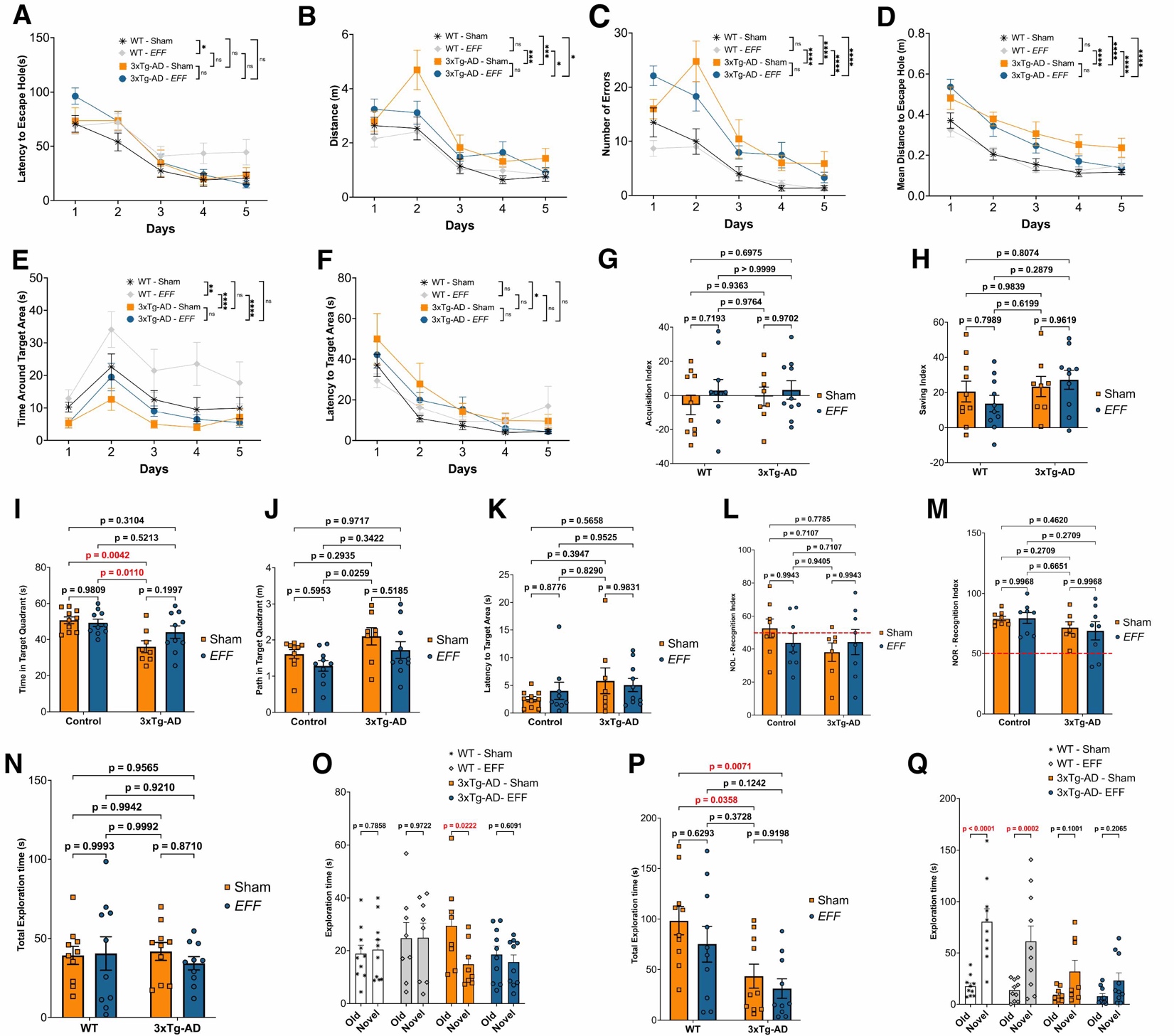

**Supplementary Figure 3. Serial injections of fibroblasts electrotransfected with *EFF* reduce cognitive capacity impairments in 3xTg-AD mice.** Evaluation of acquisition capacity of the Barnes maze spatial memory task. Several parameters are evaluated including **(A)** latency to find the escape hole, **(B)** distance traveled, **(C)** number of errors, **(D)** mean distance from the target escape hole, **(E)** time expended in the proximity of the target hole, and **(F)** latency to reach the proximity area of the target hole (n=8-10, Repeated Measures ANOVA). **(G)** Evaluation of the Barnes maze acquisition index, and **(H)** saving index (n=8-10 Two-way ANOVA). Additional parameters evaluated during the retrieval phase of the Barnes maze included **(I)** time spent in the quadrant of the target hole, **(J)** distance traveled inside the target hole quadrant, and **(K)** latency to reach the proximity area of the target hole. **(L)** Quantification of the NOL recognition index as a measure of bias exploration of a familiar object in a novel location. **(M)** Quantification of the NOR recognition index as a measure of bias exploration of a novel object. **(N)** total exploration time for the last session of the NOL task. **(O)** Quantification of the time that each mouse spends exploring the object that remained in the original location (Old) contrasted with the time spent exploring the object placed in the new location (Novel). **(P)** Total exploration time for the last session of the NOR task. **(Q)** Quantification of the time that each mouse spends exploring the familiar object (Old) contrasted with the time spent exploring the new object (Novel) (n=8-10 One-way ANOVA).

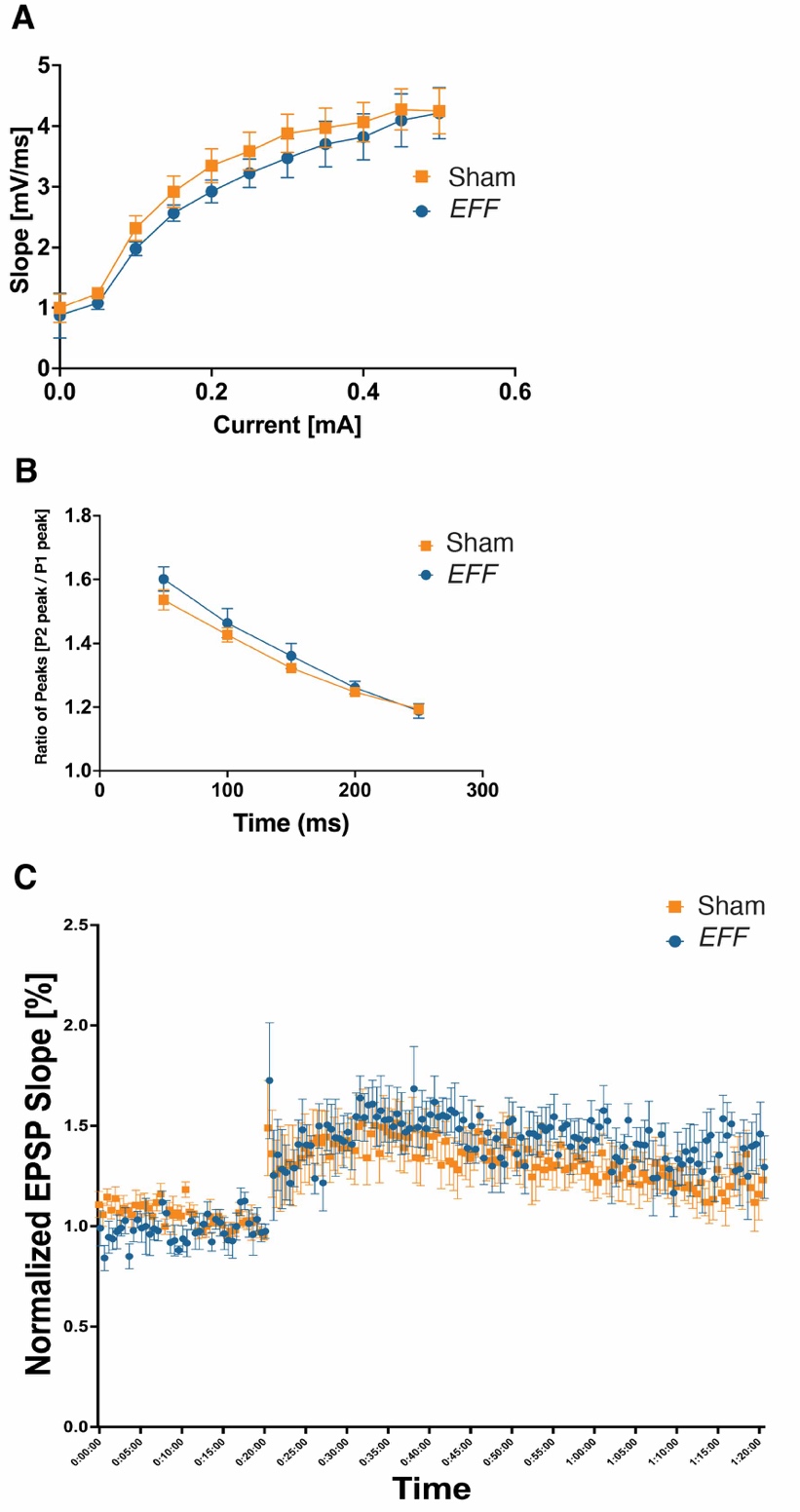

**Supplementary Figure 4. Injections *EFF-*primed cells do not induce changes in synaptic plasticity. (A)**Electrophysiological recordings of excitatory postsynaptic potential (fEPSP) from the CA1 area show no differences in the input-output (I/O) curves generated from mice injected with sham-electrotransfected cells and *EFF-*primed cells. **(B)** Paired-pulse ratio (PPR) recordings assessing transmitter release probability shows no differences in neuronal facilitation among the two groups. **(C)** Similarly, Long-Term Potentiation protocol induces similar changes in the fEPSP slope for both experimental groups.

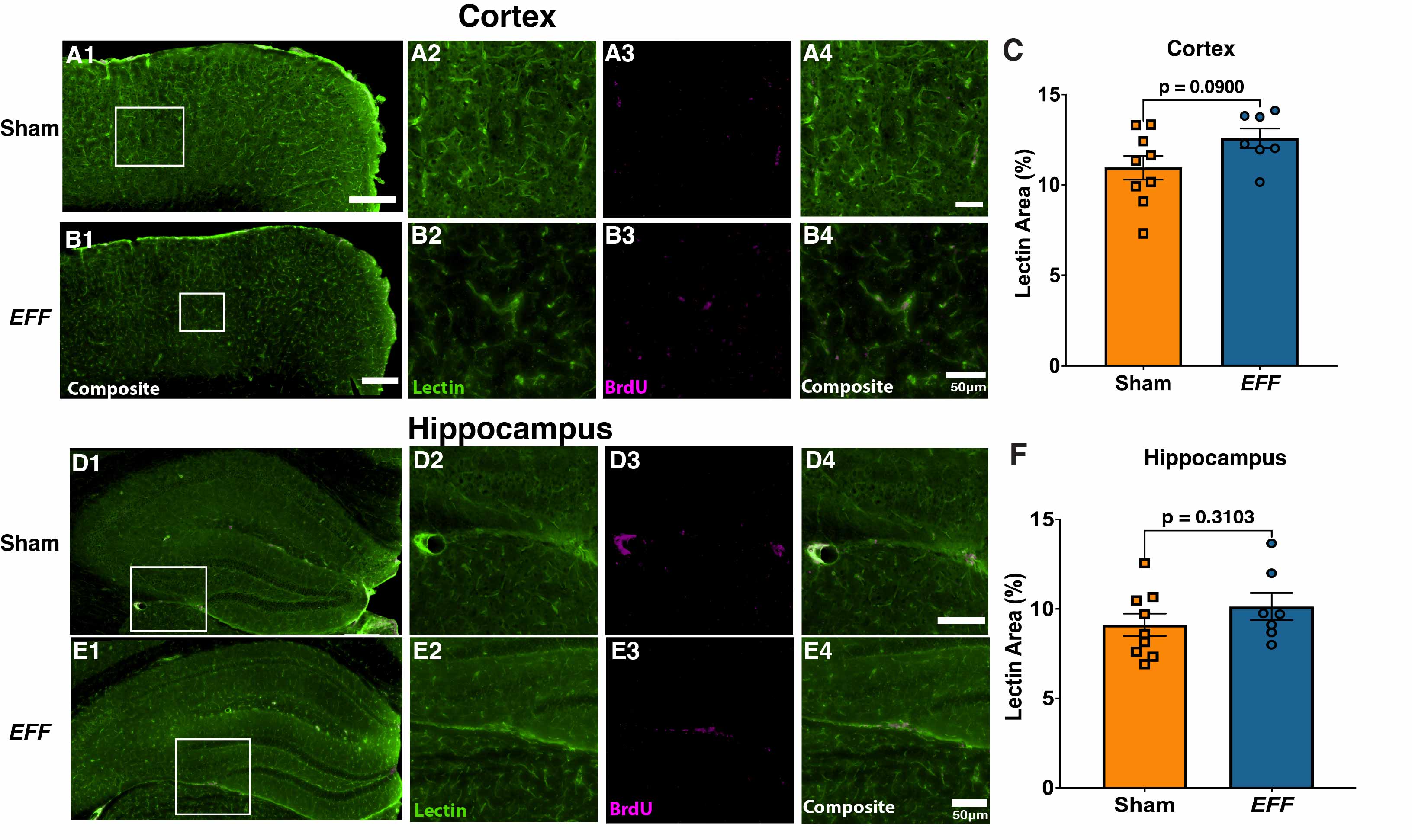

**Supplementary Figure 5. Intracranially injected *EFF-*primed cells can migrate beyond the injection site, survive long-term and increase vascularity in WT mice.** Immunofluorescence microphotographs of staining against BrdU (magenta) and blood vessels using lectin (green) in brain sections from the cortex of WT mice 6 weeks after being injected with sham-**(A1)** or *EFF*-electrotransfected cells **(B1).** **(A2-4)** Higher magnification of the area depicted in **A1**. **(B2-4)** Higher magnification of the area depicted in **B1**. **(C)** Quantification of somatosensory cortex lectin area (n= 8-10. Student’s t-test). **(D1)** Low magnification photograph of the hippocampus of WT mice injected with sham-electrotransfected cells. **(E1)** Low magnification photograph of the hippocampus of WT mice injected with *EFF*-primed cells. (**D2-4)** Higher magnification of the area depicted in **D1.**  **(E2-4)** Higher magnification of the area depicted in **E1**. **(G)** Quantification of hippocampal lectin area. (n= 8-10. Student’s t-test)

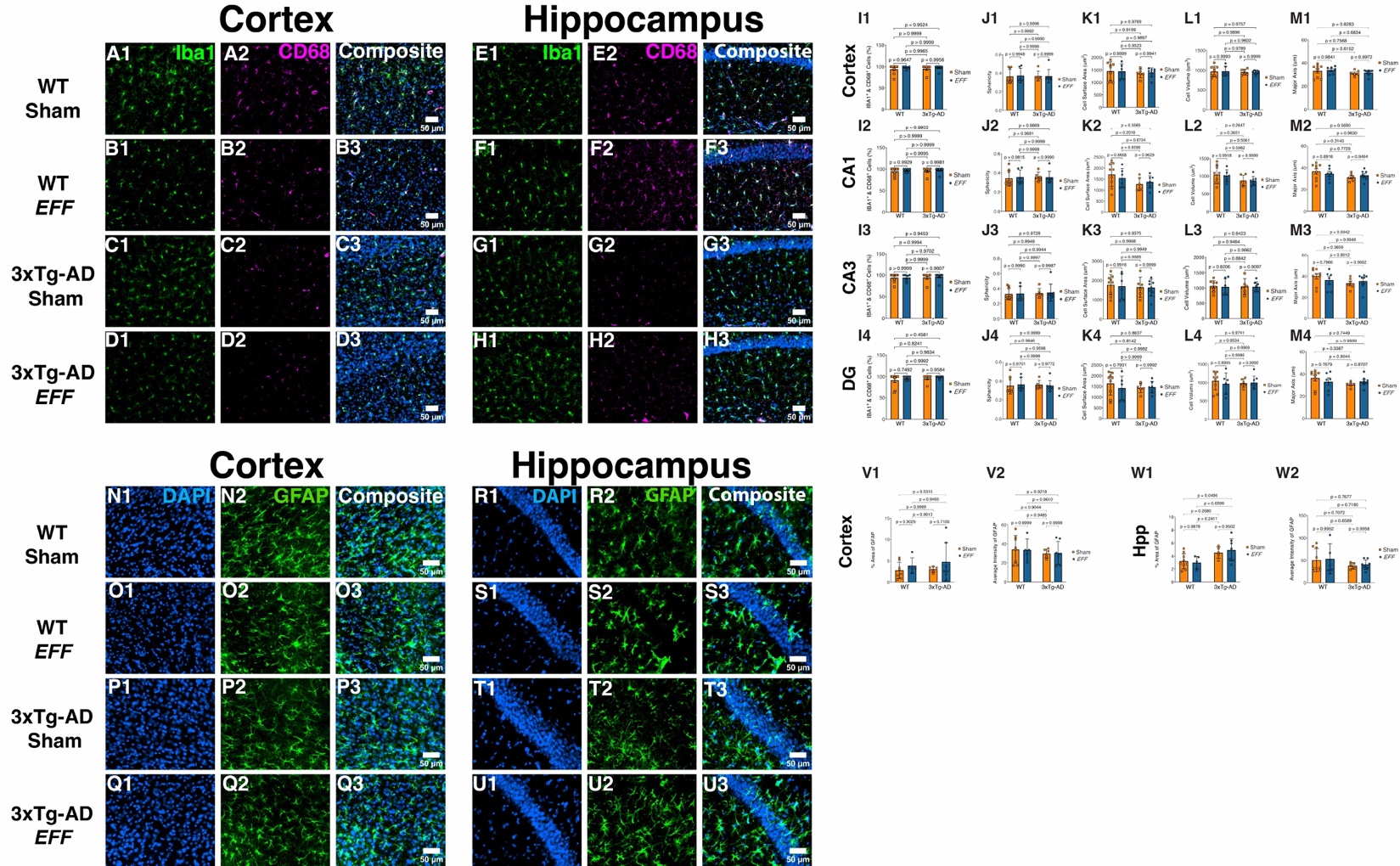

**Supplementary Figure 6. Injections of *EFF-*primed fibroblasts do not affect microglia nor astrocyte activation.** Representative microphotographs of immunofluorescent staining for the microglia marker Iba1 (green) and the microglia activation marker CD68 (magenta) in the cortex of WT mice injected with sham- **(A)** or *EFF-*primed cells **(B)**, and 3xTg-AD mice injected with sham- **(C)** or *EFF-*primed cells **(D)**. Similar representative images were taken in the hippocampus of WT mice injected with sham-**(E)** or *EFF-*primed cells **(F)**, and 3xTg-AD mice injected with sham- **(G)** or *EFF-*primed cells **(H). (I)** Quantification of percentage of activated microglia calculated by the number Iba1 cells positive for CD68 in the cortex **(I1)**, CA1 **(I2)**, CA3 **(I3)**, and **DG** (I4). In the same brain regions, parameters related to microglia activation were quantified. These parameters include **(J)** sphericity, **(K)** surface area, **(L)** cell volume and **(M)** length of the major axis. Representative microphotographs of immunofluorescent staining for the astrocyte marker GFAP in the cortex **(N-Q)** and the hippocampus **(R-U)** in control mice injected with sham cells **(N,R)** or *EFF-*primed cells **(O,S),** and 3xTg-AD injected with sham cells **(P, T)** or *EFF-*primed cells **(Q,U).** Quantification of astrocyte activation was quantified by analysis of GFAP cover area in and fluorescence intensity in the somatosensory cortex **(V)** and the hippocampus **(W).** No differences were observed between genotypes nor cell treatment (n= 8-10, Two-way ANOVA).

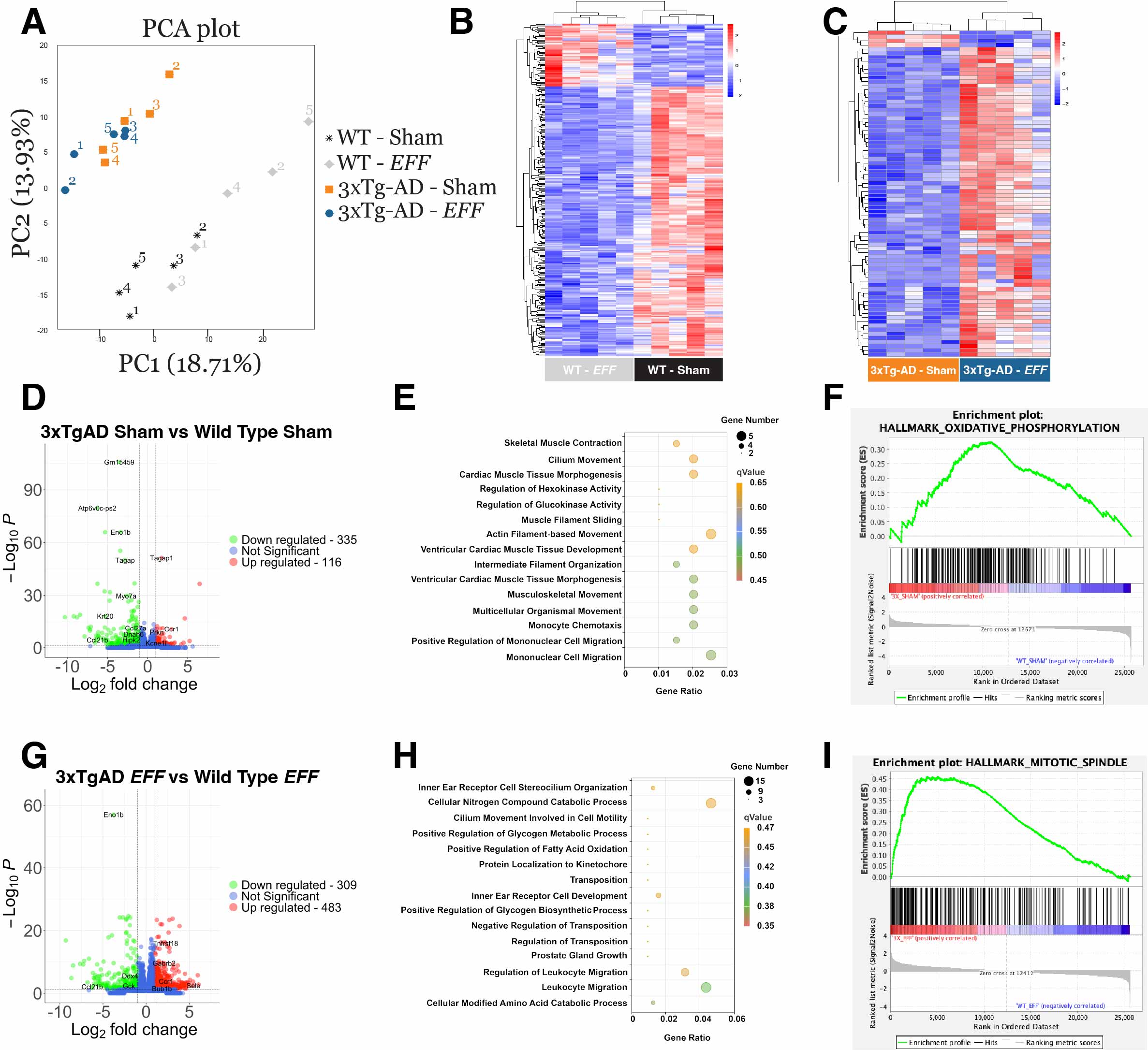

**Supplementary Figure 7. Transcriptome changes in the somatosensory cortex of WT and 3xTg-AD mice after injections of fibroblast electrotransfected with the *EFF* factors. (A)** PCA plot analysis of WT and 3xTg-AD mice injected with sham- or *EFF-*primed cells. **(B)** Unsupervised clustering heatmap of DEGs between WT mice injected with *EFF-*primed fibroblast compared with WT mice injected with sham*-*electrotransfected fibroblast. **(C)** Unsupervised clustering heatmap of DEGs between 3xTg-AD mice injected with *EFF-*primed fibroblast compared with 3xTg-AD mice injected with sham*-*electrotransfected fibroblast. **(D)** Volcano plot, **(E)** GO analysis, and **(F)** GSEA main results comparing 3xTg-AD mice and WT mice injected with Sham-electrotransfected cells. **(G)** Volcano plot, **(H)** GO analysis, and **(I)** GSEA main results comparing 3xTg-AD mice and WT mice injected with *EFF-*primed cells.

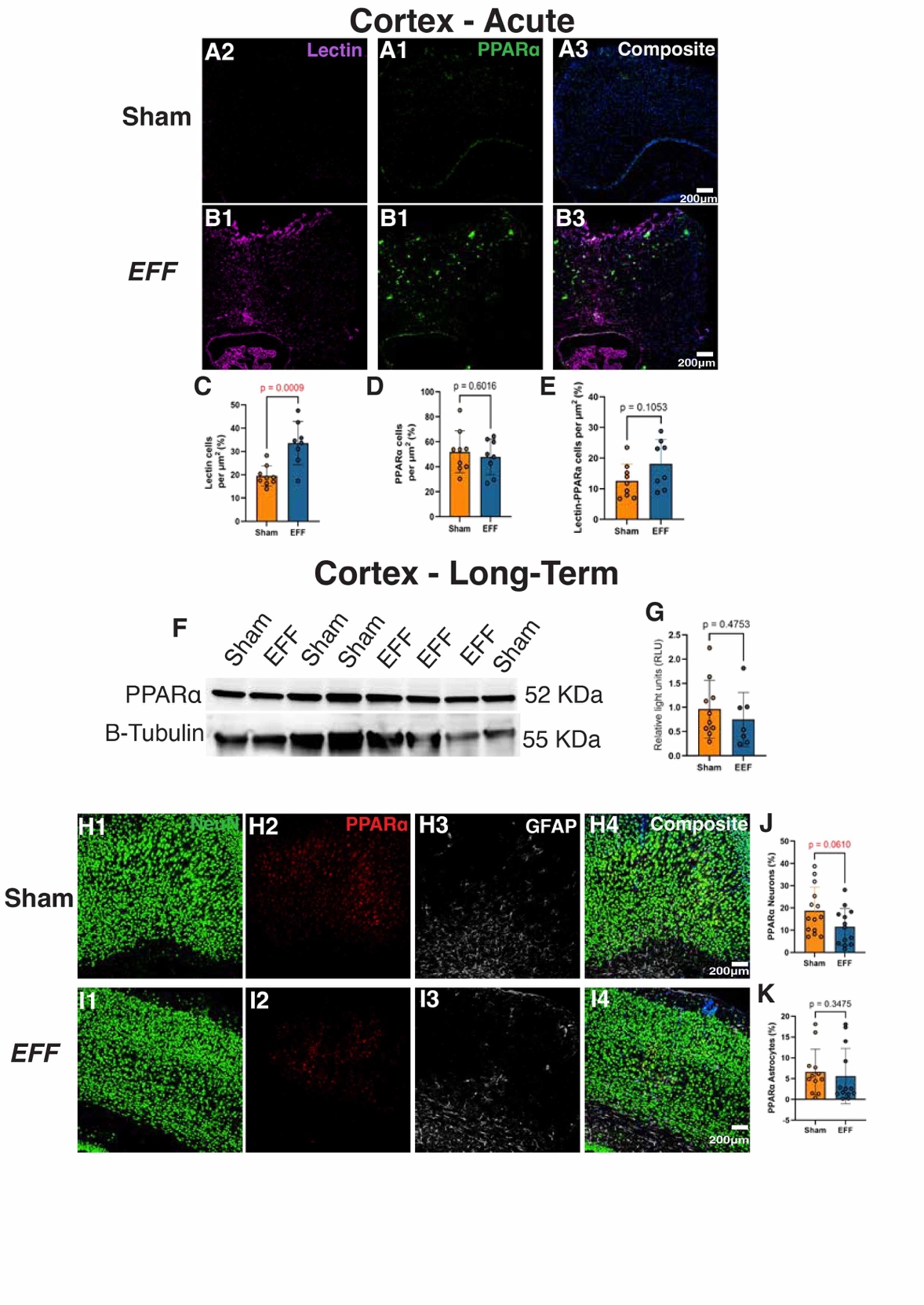

**Supplementary Figure 8. Single and multiple intracranial injections of *EFF-*primed cells do not change expression of PPARɑ in the somatosensory cortex of 3xTg-AD mice. (A,B)** Immunofluorescence staining for lectin (purple), PPARɑ (green) and DAPI (blue) in the somatosensory cortex of 3xTg-AD mice 7 days after a single injection with sham- **(A)** or *EFF-*primed cells **(B)**. Quantification of **(C)** the number of Lectin positive cells per µm^2^, **(D)** PPARɑ positive cells per µm^2^, and **(E)** PPARɑ and lectin double positive cells per µm^2^. **(F)** Western blot analysis of expression of PPARɑ protein levels and B-tubulin in the somatosensory cortex of 3xTg-AD mice 6 weeks after the last of 3 monthly injections with sham- or *EFF-*primed cells. **(G)** Quantification of relative levels of PPARɑ normalized with B-tubulin levels as housekeeping control. Immunofluorescence staining for NeuN (green) PPARɑ (red), GFAP (white) and DAPI (blue) in 3xTg-AD mice 6 weeks after the last of 3 monthly injections with sham- **(H)** or *EFF-*primed cells **(I)**. Quantification of **(J)** the percentage of NeuN and PPARɑ double positive cells, and **(K)** GFAP and PPARɑ double positive cells. (n= 8-10). Student’s t-test.

**Supplementary Tables**

**Supplementary Table 1: List of DEGs between Naïve and Sham-electrotransfected cells**

**Supplementary Table 2: List of Gene sets enriched in Sham-electrotransfected cells compared with Naïve cells**

**Supplementary Table 3: List of DEGs between Naïve and *EFF*-electrotransfected cells**

**Supplementary Table 4: List of Gene sets enriched in *EFF* -electrotransfected cells compared with Naïve cells**

**Supplementary Table 5: List of DEGs between Sham- and *EFF*-electrotransfected cells**

**Supplementary Table 6: List of Gene sets enriched in *EFF* -electrotransfected cells compared with Sham-electrotransfected cells**

**Supplementary Table 7: List of DEGs in WT mice injected with Sham- or *EFF*- electrotransfected cells**

**Supplementary Table 8: List of Gene sets enriched in WT mice injected with Sham- electrotransfected cells compared with WT mice injected with *EFF*- electrotransfected cells.**

**Supplementary Table 9 : List of DEGs in 3xTg-AD mice injected with Sham- or *EFF*- electrotransfected cells**

**Supplementary Table 10: List of Gene sets enriched in 3xTg-AD mice injected with Sham- electrotransfected cells compared with 3xTg-AD mice injected with *EFF*- electrotransfected cells**

**Supplementary Table11: List of Plasmids used for electrotransfection**

**Supplementary Table 12: List of Primers used for qRT-PCR analysis**

**Supplementary Table 13: List of Primary and Secondary antibodies**
